# Differential aggregation patterns of *Endozoicomonas* within tissues of the coral *Acropora loripes*

**DOI:** 10.1101/2024.12.17.629048

**Authors:** Cecilie R Gotze, Ashley M Dungan, Allison van de Meene, Katarina Damjanovic, Gayle K Philip, Justin Maire, Lone Hoj, Linda L Blackall, Madeleine JH van Oppen

**Affiliations:** School of BioSciences, The University of Melbourne, Parkville, VIC 3010, Australia; Australian Institute of Marine Science, Townsville, QLD 4810, Australia; Ian Holmes Imaging Centre, Bio21, The University of Melbourne, Parkville, VIC 3010, Australia; Melbourne Bioinformatics, The University of Melbourne, Parkville, VIC 3010, Australia

**Keywords:** *Endozoicomonas*, coral symbiosis, microbiome, symbiotic bacteria

## Abstract

Bacteria in the genus *Endozoicomonas* are well-known coral symbionts commonly found as clusters within tissues of several coral species. Mapping the spatial distribution of these microbial communities is critical to gaining a holistic understanding of the potential role they may play within the coral host. This study focuses on characterising bacterial aggregates associated with the common reef-building coral, *Acropora loripes*, from the central Great Barrier Reef, Australia. A conventional cultivation-based method was employed to establish a pure culture collection of 11 undescribed *Endozoicomonas* strains isolated from *A. loripes*. Subsequent 16S rRNA gene sequence analysis revealed their classification into two distinct phylogenetic clades. To resolve their spatial distribution *in hospite*, clade-specific fluorescence *in situ* hybridisation probes were designed. Aggregates were consistently observed in the gastrodermal tissue layers surrounding the upper and lower gastrovascular cavity and were predominantly formed by cells from the same phylogenetic clade, with a minor proportion of aggregates formed by *Endozoicomonas* from both targeted clades. Furthermore, a clear distinction in aggregation pattern was observed; one clade exhibited clusters with regular and contained growth patterns, whereas the other formed clusters lacking clear boundaries and having irregular shapes. Scanning electron microscopy revealed the presence of a membrane of unknown origin associated with bacterial aggregates in two instances, suggesting potential structural or functional differences in these aggregates. These contrasting morphological features underscore the need for comprehensive investigations into the underlying mechanisms governing bacterial aggregate formation in corals.

## Introduction

Coral reefs are known as the rain forests of the sea, as they are among the world’s most biologically diverse ecosystems (1). Much of the ecological success of reef-building corals is attributed to their endosymbiosis with dinoflagellate algae in the family Symbiodiniaceae (2,3). However, corals also form associations with a multitude of other microbes, including other protists, bacteria, fungi, archaea, and viruses (4–7) Together, the coral host and its associated microbial community form a multipartite organism termed a holobiont. Historically, coral microbiology has focused on the coral-Symbiodiniaceae symbiosis, but the last decade has seen increasing recognition of the role bacteria play in holobiont functioning and health (8–10). However, studies aimed at deciphering the roles of bacteria within the holobiont are complicated by the diverse and rich nature of their bacterial communities, with some coral species harbouring several thousand different prokaryotic taxa (11). Additionally, coral-associated bacterial communities have also been demonstrated to differ according to environmental disturbances, seasonality, geographic area, coral genotype, and even within a single colony (12–16).

A reductionist approach can be employed by studying coral species that naturally host relatively low-diversity bacterial communities, with some corals naturally associating with as few as 10-20 bacterial species, often dominated by a few highly abundant taxa (17–20). Among these, the bacterial genus *Endozoicomonas* has garnered significant interest due to its association with various marine invertebrates, such as cnidarians (21,22). The global prevalence and abundance of *Endozoicomonas* within coral ecosystems suggest a pivotal ecological role, potentially driven by competitive advantages or mutualistic relationships with the coral hosts (23,24). Although the functional role of *Endozoicomonas* in the host microenvironment is not fully understood, they are suggested to be involved in fundamental holobiont processes such as B-vitamin metabolism (25) as well as cycling of nitrogen, sulfur, and phosphorus (26–28).

The generally high relative abundance of *Endozoicomonas* within corals can be attributed to their capacity to form dense clusters, known as cell-associated microbial aggregates (CAMAs). While CAMAS have been found in a diversity of coral species throughout the Indo-Pacific (29) and the Caribbean (30,31), the presence of *Endozoicomonas* in these aggregates has only been confirmed in a subset of coral species. This includes CAMAs occurring within the epidermal layer of the tentacles of corals from the family *Pocilloporidae* (28,32,33). Moreover, members of the novel genus, *Soroendozoicomonas*, within the Endozoicomonadaceae family, were recently identified as forming aggregates in the mesenterial filaments in the coral *Pocillopora acuta*, which are used for digestion and prey capture (34). The lack of a consistent pattern in aggregate distribution suggests that the process may be phylotype-specific, host-specific, or influenced by other host physiological factors. However, the microorganisms that reside within these aggregates have not been fully characterised.

Only one recent study has applied species-specific probes to investigate the bacterial composition within CAMAs and found them to be occupied by multiple *Endozoicomonas* phylotypes, indicating the potential for shared metabolic pathways (28). Nonetheless, most studies of *Endozoicomonas* have either described their abundance or genomic repertoire without assessing their spatial distribution and ability to form CAMAs (25,27,35,36). Consequently, important knowledge on the taxonomic composition of bacteria within CAMAs and how they are distributed throughout different microhabitats of the host is lacking. Such knowledge could help decipher their functional potential, as different locations likely represent distinct metabolic and biochemical niches where members could carry out different functions (37).

To address this knowledge gap, we targeted the common reef-building coral *Acropora loripes,* reported to have low-diversity bacterial communities (19,20). We employed a culture-based approach combined with fluorescence *in situ* hybridisation (FISH) and scanning electron microscopy (SEM) to visualise the spatial distribution, taxonomic composition, and morphology of CAMAs within their native host, obtained from two reef sites on the central Great Barrier Reef (GBR), Australia., i.e., Davies and Backnumbers Reef.

## Materials and methods

### Coral Collection and Maintenance

Visually healthy *A. loripes* colonies (25 cm diameter) were collected from two sites on the central Great Barrier Reef: Davies Reef (June 2020, February 2021) and Back Numbers Reef (December 2020) (Permit G12/35236.1). Corals were transported to the Australian Institute of Marine Science (AIMS) in aerated seawater and maintained at the National Sea Simulator (SeaSim) under ambient lighting and temperature conditions that mirrored the Davies Reef profile. Corals were supplied with 0.4 µm filtered seawater at a turnover rate of 4.8 times per day and fed daily with *Artemia* (0.5 nauplii mL⁻¹). Additional collection details and tank parameters are provided in Supplementary Materials.

### 16S rRNA Gene and ITS2 Metabarcoding

To characterise the bacterial communities associated with *A. loripes*, fragments were collected from four distinct locations within each coral colony to capture intra-colony microbial diversity. DNA was extracted from 3–5 mm fragments, followed by amplification of the V5– V6 region of the 16S rRNA gene and the ITS2 region of Symbiodiniaceae. Sequencing was performed using Illumina technology at the Walter and Eliza Hall Institute (Melbourne, Australia). Amplicon data were processed using QIIME2, and ITS2 sequences were analysed with the SymPortal framework. For additional details, including primer sequences, PCR protocols, and bioinformatics pipelines, see Supplementary Materials.

### Bacterial Culturing

Coral fragments were processed to isolate tissue-associated bacteria using tissue homogenization, followed by serial dilutions and plating on Marine Agar 2216. Plates were incubated in the dark at 23°C, and bacterial colonies were purified over three passages. DNA from bacterial isolates was screened using PCR with *Endozoicomonas*-specific primers and sequenced to confirm identity. Further details, including homogenization protocols, culturing conditions, and PCR reagents, are provided in Supplementary Materials.

### Phylogenetic Analysis

Phylogenetic relationships of cultured *Endozoicomonas* strains were assessed by constructing a maximum likelihood tree of 16S rRNA sequences aligned with near full-length Endozoicomonadaceae sequences from the SILVA database. Sequences were aligned using MAFFT, and the TIM3e+R6 model was selected with ModelFinder. Bootstrap support for tree topology was evaluated with 1000 replicates. Detailed pipeline steps, including software versions and alignment methods, are available in Supplementary Materials.

### Design and Validation of FISH Probes

Phylotype-specific probes for *Endozoicomonas* Clade-A and Clade-B were designed using the ARB probe design tool with a curated database of 16S rRNA sequences. Probes were tested for specificity using in silico validation and hybridization with pure bacterial isolates. Stringency conditions were optimized using increasing formamide gradients (15–30%). See Supplementary Materials for probe sequences, specificity testing, and optimisation.

### Sample Fixation, Histology, and microscopy

Coral fragments were fixed in 4% paraformaldehyde, decalcified in EDTA, and embedded in paraffin for histology and FISH. Tissue sections were stained with hematoxylin and eosin or hybridized with clade-specific and universal *Endozoicomonas* probes labeled with Atto550 or Atto647N. Autofluorescence was reduced through methanol quenching, and probe specificity was confirmed via confocal laser scanning microscopy. For SEM, paraformaldehyde-fixed tissue sections where CAMA were identified were further processed with heavy metal staining (uranyl acetate and lead citrate) to enhance contrast and the CAMA visualized using a scanning electron microscope after correlation with the optical image. Detailed fixation, staining, and imaging protocols are described in supplementary materials.

## Results

### Microbial community composition within the tissues of the coral *A. loripes*

To characterise the bacterial community composition associated with *A. loripes*, metabarcoding analysis of the 16S rRNA gene (V5-V6 region) was conducted on coral fragments from 12 of the 14 coral colonies collected from the GBR, Australia. Bulk tissue samples (n=4 per coral colony) indicated a similar microbial community composition across all sampled colonies, comprising just a few bacterial genera (Fig. 1A). Furthermore, considering all amplicon sequence variants (ASVs), including rare ones, a total of 600 ASVs were identified from 47 samples across 12 *A. loripes* corals. However, upon excluding low-abundance ASVs (i.e., those with counts representing less than 0.001% relative abundance), a total of 249 ASVs were present across all 47 samples. Notably, *Endozoicomonas* was found as the dominant genus across almost all replicate samples, with sequences affiliated with *Endozoicomonas* accounting for more than 95% of the relative abundance (RA) in some samples. Other major taxa included *Candidatus amoebophilus* (colony H, ∼25% RA), *P3OB-42* (family Myxococcaceae, in coral Al06, ∼15% RA), and *Candidatus Fritschea* (family Simkaniaceae, coral Al01, ∼10% RA) and an unknown taxon in of Gammaproteobacteria (coral Al06 and Al09, ∼20% RA).

**Fig. 1.**
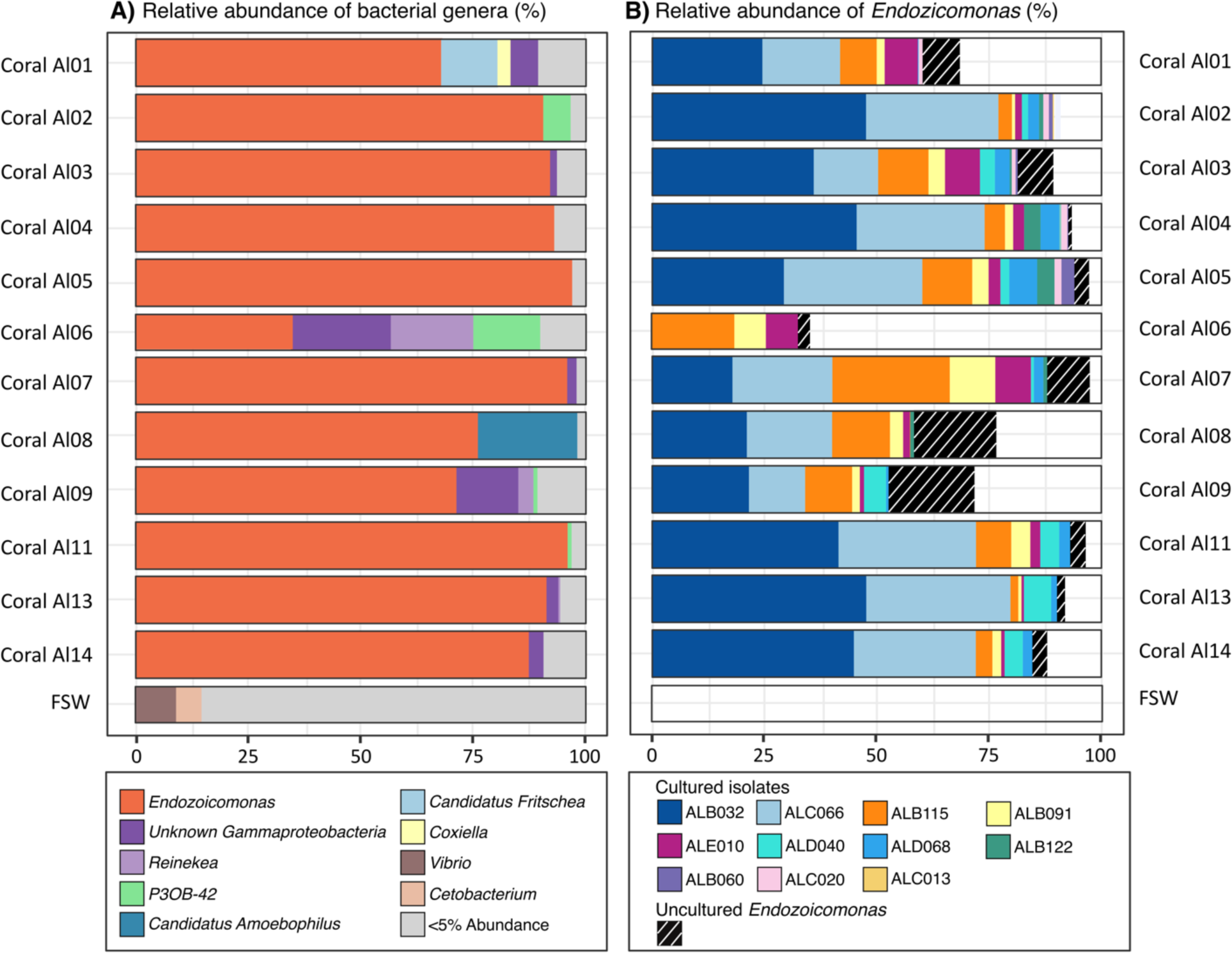
Relative abundance of *A. loripes*-associated bacteria. **A** Bar plot visualising the relative abundance of bacterial genera identified in March 2021 across 12 *A. loripes* colonies annotated on the vertical axis. Genera with <5% relative abundance were pooled into the “<5% Abundance” category. Each coral colony (Al01-Al14) was subsampled across four distinct-colony locations, with replicate samples pooled by genotype for subsequent analysis. Relative abundances are represented as the average across four replicate samples. Three replicate samples of filtered seawater from the aquarium where the corals were reared were included in 16S rRNA gene metabarcoding for comparison (FSW). **B** Bar plot showing the relative abundance of the ASVs matching the cultured or uncultured strains across the same 12 *A. loripes* colonies. Relative abundances are represented as the average across the four replicates and two time points (March and June 2021). The hatched proportion represents ASV’s which did not match any isolate sanger sequences, i.e. these were not obtained in the *Endozoicomonas* culture collection.

In addition to 16S rRNA gene community composition analysis, internal transcribed spacer 2 (ITS2) metabarcoding was carried out to characterise the endosymbiotic microalgal taxonomic composition. Analysis of ITS2 amplicon data obtained from bulk tissue samples (n=4 per coral colony) revealed that all coral colonies hosted a single Symbiodiniaceae genus, *Cladocopium*. While pairwise adonis comparisons of ITS2-type profiles indicated compositional differences in some coral colonies (*p* = 0.005), this had no significant effect on the microbiome composition (Fig. S6).

### Culturing the dominant taxa from the microbiome of *A. loripes*

At the same time as the metabarcoding samples were taken, *A. loripes* colonies were sampled to establish pure cultures of associated bacteria on Marine Agar. Culturing efforts yielded >500 bacterial pure cultures, among which 87 were identified as belonging to *Endozoicomonas* through PCR screening with genus-specific primers. Analysis of the 16S rRNA gene sequences from each isolate revealed 11 distinct *Endozoicomonas* strains. The comparison of isolate-derived near full length 16S rRNA gene sequences with ASV sequences from the metabarcoding data of the coral holobiont demonstrated that the culture collection represented the dominant ASVs associated with *A. loripes,* with the culturable proportion of *Endozoicomonas* ASV’s ranging from 28% in coral Al06 to 95% in coral Al05 (Fig. 1B). Among the isolated bacteria, three *Endozoicomonas* isolates, namely ALB032, ALC066, and ALB115, were the most abundant in *A. loripes* based on matches to ASVs in the metabarcoding dataset. However, their relative abundance exhibited marked variations across coral colonies, with ALB115 showing low relative abundance (<1%) in colony Al13.

### Phylogenetic distinction of newly isolated *Endozoicomonas* strains

To evaluate the taxonomy and phylogenetic connections among newly isolated *Endozoicomonas* strains, we constructed a maximum likelihood phylogeny using near-complete 16S rRNA sequences of. Alongside our novel isolates, we incorporated 1459 Endozoicomonadaceae 16S rRNA sequences >1200 bp sourced from the SILVA database. The four isolates ALB091, ALE010, ALB115, and ALC013 share >97% sequence identity in their 16S rRNA gene, indicating that they belong to the same species (De Albuquerque and Haag 2023)(Fig. 5), hereafter referred to as Clade-A. Isolates ALB060, ALC020, ALC066, ALD068, ALB122, ALB032 and ALB040 also exhibited >97% sequence identity and together formed a distinct clade, named Clade-B. Pair-wise comparisons between 16S rRNA gene sequences from each clade revealed that Clade-A and Clade-B on average share 94% identity. Moreover, phylogenetic analysis (Fig. 2) indicated that despite low sequence identity, they both share common recent ancestors with *Endozoicomonas* sequences obtained from the coral *Acropora humilis*, sampled from the Red Sea (38,39).

**Fig. 2.**
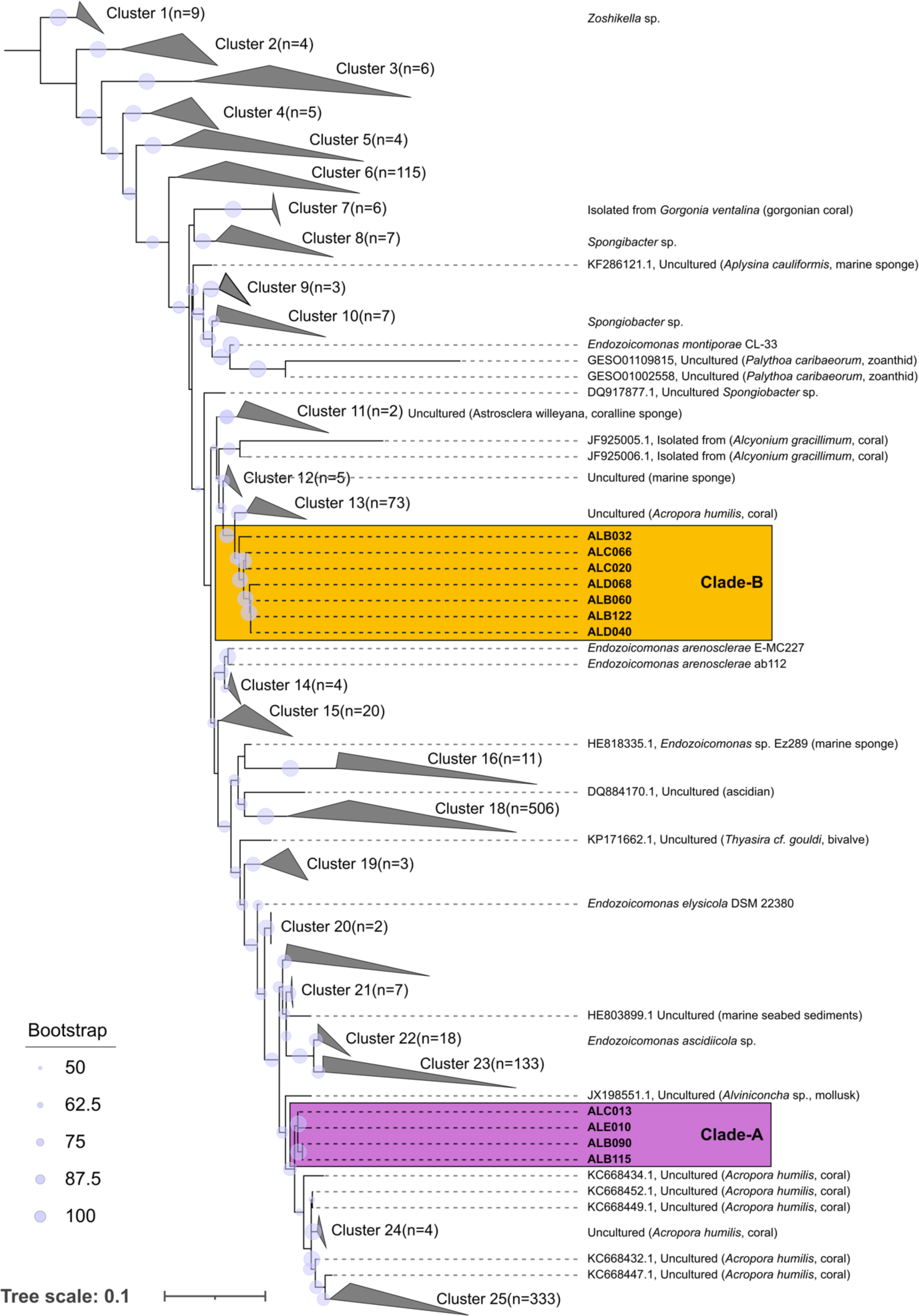
Phylogenetic placement of isolated *Endozoicomonas* strains. Placement of 11 *A. loripes*-associated *Endozoicomonas* strains within the Endozoicomonadaceae family. Maximum likelihood tree constructed from 1459 bacterial 16S rRNA gene sequences (length of total alignment 2087bp) obtained from the SILVA database in addition to 16S rRNA gene sequences extracted from 11 novel *A. loripes* sourced isolates. Bootstrap values greater than 50% based on 1000 replications are shown at the nodes. The tree is rooted at *Zoshikella*.

### Stability and composition of *Endozoicomonas* after three months in captivity

To assess the stability of the bacterial community composition, coral colonies sampled in March 2021 were re-sampled in June 2021. During this period, corals were kept in an aquarium under conditions mimicking seasonal ambient light and temperature at Davies Reef, GBR. Replicate samples from the same coral colony were pooled for each time point. Metabarcoding analysis revealed that *Endozoicomonas* remained the dominant taxon across all coral colonies (except Al06, which was excluded early due to tissue sloughing) (Fig. 3). While some variation in the relative abundance of specific *Endozoicomonas* amplicon sequence variants (ASVs) was observed, the overall prevalence of *Endozoicomonas* persisted over time.

**Fig. 3.**
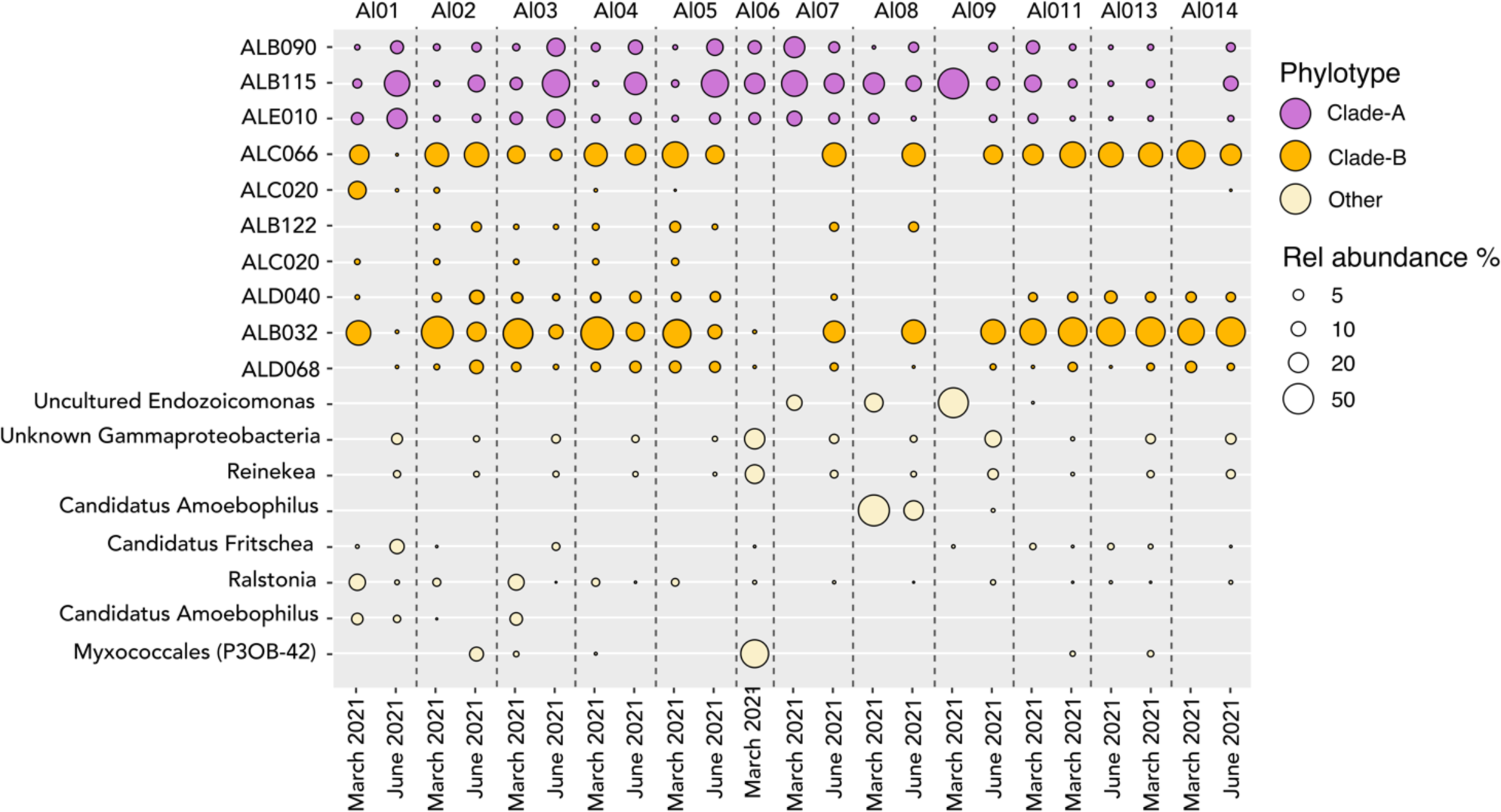
Relative abundance across two timepoints. Relative abundance of the 18 most abundant 16S rRNA amplicon sequence variants (ASVs) over time in captive *A. loripes.* Bubble plot showing the relative abundance (%) of the 18 most abundant ASVs across all 12 *A. loripes* colonies. Each column visualises ASVs in one representative colony annotated above (Al01-Al14). Relative abundances are represented as the average across four replicate samples. Sampling for 16S rRNA gene metabarcoding analysis was carried out in March and June 2021.

### *In situ* spatial distribution and composition of CAMAs

To examine the spatial distribution of the two major phylotypes (Clade-A and Clade-B) within the host tissue, coral colonies sampled for 16S rRNA gene metabarcoding analysis in June 2021, were concurrently sampled for histological examination. Six replicate fragments (∼2 cm branch tips) were sampled from different locations across each coral colony ranging from sun-exposed to shaded areas. A total of 293 CAMAs were identified by H&E staining of the serial-sectioned tissues derived from distinct colonies. To validate the presence of *Endozoicomonas* within the CAMAs, we also conducted FISH with previously published probes targeting either all bacteria (40) or specifically the genus *Endozoicomonas* (38)(Fig. 4). By employing both probe types on consecutive serial sections, we confirmed that *Endozoicomonas* constitute the primary and potentially sole, bacterial genus forming CAMAs within *A. loripes*.

**Fig. 4.**
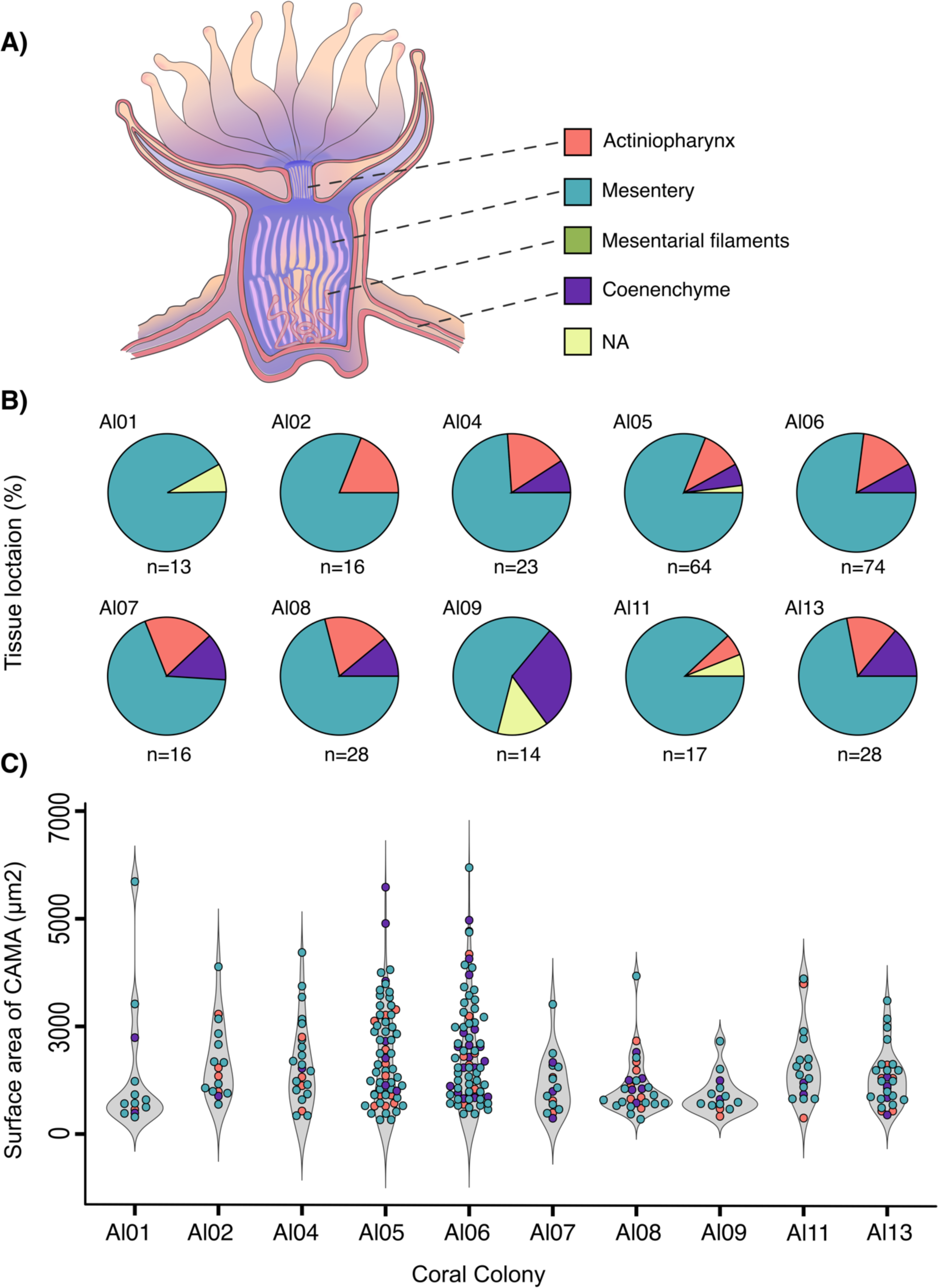
CAMA distribution and size proxy across anatomical regions. 2D surface area measurements of 4 μm coral tissue sections showing CAMAs within various tissues of *Acropora loripes*. **A** Illustration of a coral polyp with color-coded tissue regions where CAMAs were detected (refer to panels B and C). **B** Pie charts showing the frequency of CAMA occurrence in the various tissue compartments. **B** Dot plots comparing CAMA sizes (surface area in 2D sections), with colors representing the tissue compartment of each CAMA. A total of 293 CAMAs were detected in H&E-stained tissues. Violin plots show the median and interquartile range of CAMA sizes from each coral colony. “NA” denotes CAMAs with indeterminate locations due to histological artifacts. Schematic representation inspired by (41)

*Endozoicomonas* CAMAs were found distributed extensively across several regions of the coral polyp, predominantly associated with the gastrodermal layer lining the mesenteries, which forms multiple folds that span the upper and lower gastrovascular cavity (Fig. 4)(42). In total, 78% of CAMAs were detected in the mesenterial gastrodermis, 13% in the gastrodermis lining the actinopharynx, and 7% in the coenenchyme, which acts as gastrovascular tubes between polyps. The precise location of the remaining 2% could not be determined due to histological artefacts. On average three CAMAs were detected per 2 cm longitudinal coral section. Exceptions were corals Al01 and Al07, in which histological sections appeared not to have been cut deep enough to capture tissues lining the gastrovascular cavity. The average diameter of CAMAs across all corals was 42.8 µm (SD = 17.5, SE = 1.02), with the largest being 97 µm and the smallest 12 µm - a ninefold difference. This considerable variation may, in part, result from the specific plane of each CAMA captured in the 4 µm-thick histological sections.

### Clade-Specific Aggregation and Morphology of CAMAs

To further resolve the composition and spatial distribution of *Endozoicomonas* within CAMAs, we employed FISH with newly designed clade-specific oligonucleotide probes targeting either Clade-A or Clade-B (Fig. 3). Each coral colony underwent an investigation of 30 CAMAs to discern the bacterial composition at the clade level based on observed hybridisation patterns. The analysis revealed that the majority of CAMAs consisted of members belonging to Clade-B, comprising 83% of CAMAs observed with phylotype-specific FISH probes. These Clade-B CAMAs were present across all examined coral colonies. In contrast, CAMAs composed of Clade-A members were exclusively identified in corals Al01, Al05, Al06, Al07, and Al11.

A distinct morphological pattern differentiated the two clades: CAMAs formed by Clade-A displayed a regular and contained aggregation pattern, whereas those formed by Clade-B lacked a clear boundary, exhibiting irregular shapes (Fig. 5). While Clade-B CAMA morphologies ranged from highly dispersed growth to somewhat more circular shapes, they never appeared to be enclosed by a restricted boundary in the outer periphery as consistently observed in Clade-A CAMA morphologies (Fig. S8). To resolve the clade-specific CAMA patterns, we examined sectioned coral tissue using SEM. In coral genotype Al07, this analysis revealed bacterial aggregates enclosed by a distinct membrane of unknown origin (Fig. 8). However, no SEM images of Clade-B CAMAs were obtained, as Clade-B aggregates could not be detected in the examined tissue sections.

**Fig. 5.**
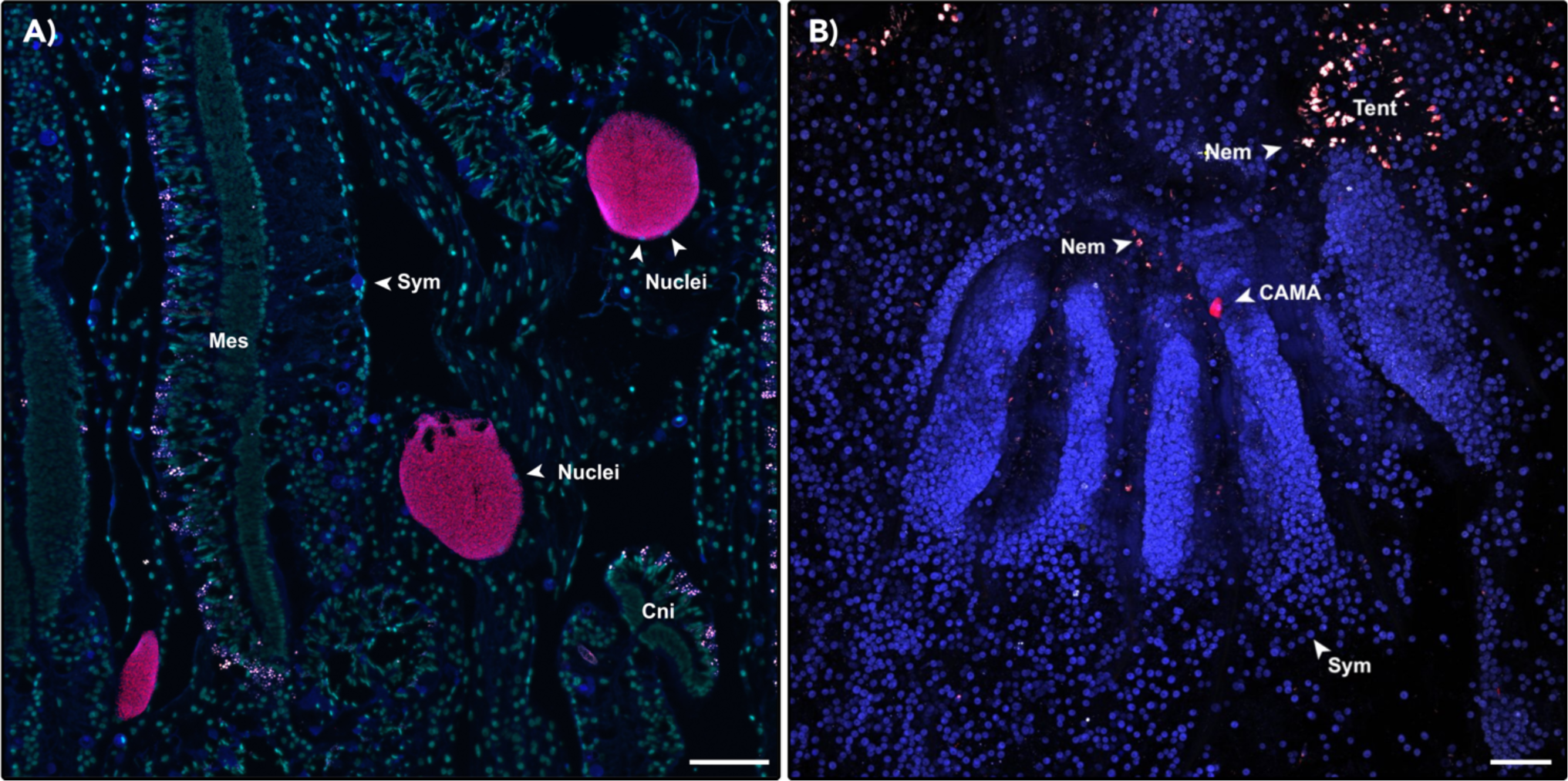
CAMAs found within the tissue of *A. loripes*. **A** CAMAs displaying hybridization with the ‘All-*Endozoicomonas* mix’ probe (red) targeting all *Endozoicomonas* bacteria are localised within the mesenteries lining the gastrovascular cavity. **B** Whole-mount FISH image showing a CAMA in the actinopharynx hybridised with the ‘All-*Endozoicomonas* mix’ probe (red). In both images, red fluorescence corresponds to the ‘All-*Endozoicomonas* mix’ probe labelled with Atto647N. Non-EUB probe labelled with Atto550 (negative control) is depicted in white. Cyan indicates DNA stained by DAPI (shown in **A** only). Host autofluorescence is represented in blue while overlapping signals between the target probe and negative control probe are shown in pink indicating non-specific binding at nematocysts and mucocytes within the Cnidoglandular bands. Sym, Symbiodiniaceae; Nuclei, Host nuclei; Nem, Nematocytes; Tent, tentacle; Cni, Cnidoglandular bands. Scale bars represent 100 μm.

Moreover, CAMAs where both clade-specific oligonucleotide probes hybridised with cells in the same CAMA, were only observed six times in corals Al07, Al08, and Al09. The morphology of all six CAMAs which were comprised of both *Endozoicomonas* clades closely resembled that of Clade-A (Fig. 5A). In some mixed CAMAs, fluorescent signals from the Clade-A specific probe were concentrated at the outer edges, while in others, both probes exhibited a homogeneous distribution across the same CAMA (Fig. 6). However, no clear patterns were identified regarding the cellular arrangement of Clade-A and Clade-B within mixed-population aggregates. Lastly, we found no CAMAs that did not bind phylotype-specific oligonucleotide probes, emphasising that CAMAs were exclusively formed by members belonging to either Clade-A, Clade-B, or a combination of both.

**Fig. 6.**
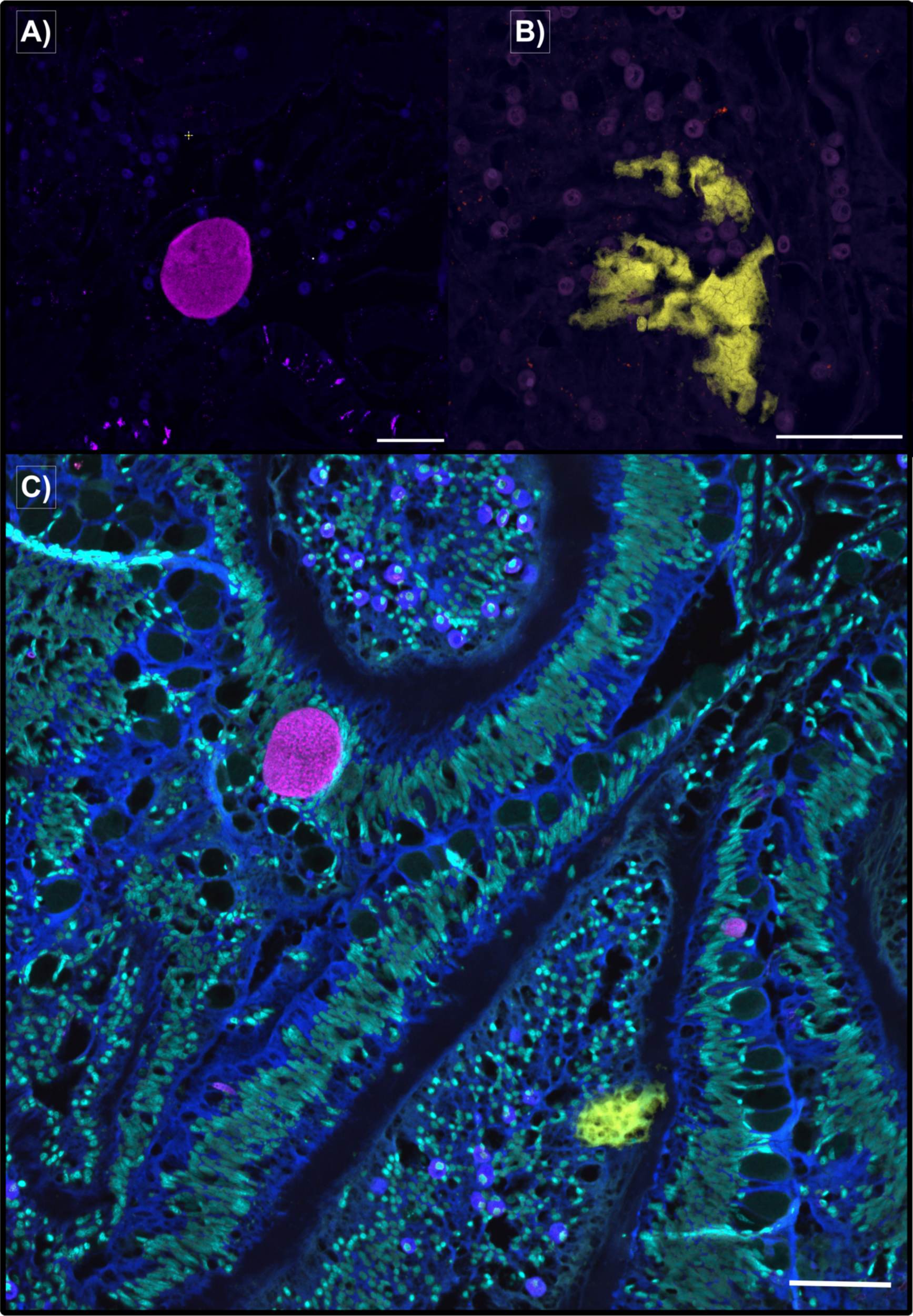
Clade-specific CAMA morphology. **A** CAMA observed in a tissue section, showing hybridisation signal from the Clade-A specific probe labelled with Atto647N depicted (depicted in magenta). **B** CAMA in a tissue section hybridised with the Clade-B specific probe labelled with Atto550 depicted in yellow. **C** Three adjacent CAMAs, with two CAMAs hybridized with the Clade-A probe (magenta) and another CAMA hybridized with the Clade-B probe (yellow). Blue indicates host autofluorescence, while cyan highlights nuclear structures stained by DAPI. Scale bars represent 50 μm.

## Discussion

### Dominant *Endozoicomonas* species of *A. loripes* are stable symbionts

Overall, the relative abundance of *Endozoicomonas* within captive *A. loripes* colonies persisted over three months, which is consistent with previous findings in *A. loripes* (19), as well as in other coral families (32,43). A consistently high relative abundance of three ASVs was observed, i.e. ALB032, ALC066 from Clade-B and ALB115 from Clade-A, and suggests these isolates could be true symbionts of *A. loripes* forming a temporally stable association with their coral host. However, genotype-specific differences in the relative abundance of ALB032, ALC066, and ALB115 were observed, and the distribution of these ASVs varied across individual corals (Fig. 3). However, it is important to note that this study examined relative abundance only, and fluctuations may also be influenced by the presence of large bacterial aggregates within sampled fragments causing bacterial abundances to increase (44,45). Therefore, to draw more conclusive insights, ASV stability should ideally be assessed using absolute abundance data (20,46).

### Diversity and co-occurrence of *Endozoicomonas* within *A. loripes*

Phylogenetic analysis (Fig. 2), along with their relatively low sequence identity, indicated that *Endozoicomonas* strains isolated from tissues of the coral *A. loripes* belonged to two distinct species (47) co-occurring within the same host. However, not all *Endozoicomonas* ASVs were detected in all sampled colonies, for instance, ALB112 and ALC020 from Clade-B were only detected in a subset of corals, entirely lacking in 7 out of the 12 corals sampled (Fig. 3). A similar pattern has been observed in other *Acropora* species, where multiple distinct *Endozoicomonas* phylotypes were obtained from the same corals, although with widely varying abundance among individual corals (35,39). This pattern is not only restricted to acroporid corals but has also been observed in pocilloporids (48), suggesting that a subset of strains in this study could be more widespread, possibly originating from differing adaptative strategies despite occurring within the same coral host (10,25). These strategies could include variations in resource acquisition mechanisms, metabolic flexibility, or tolerance to specific environmental stressors, allowing distinct *Endozoicomonas* strains to coexist while occupying different ecological niches of the same coral host (23). Moreover, local environmental conditions may drive associations with specialised *Endozoicomonas* phylotypes. For instance, corals in the genus *Pocillopora* associate with *Endozoicomonas* phylotypes that are globally distributed, whereas *Stylophora* exhibits a higher degree of host-microbe specificity where colonies associate with unique or closely related *Endozoicomonas* strains (33,36,38). Thus, some corals might select specific characteristics or functions that different *Endozoicomonas* strains and species have and which are required for adaptation to their particular ecological niche or metabolic requirements (49). Conversely, instead of the host-selection of bacterial symbionts, it is plausible that the bacteria colonise hosts based on their nutritional requirements which may be provided by coral or photosymbiont, as some coral-associated bacteria appear to be correlated with the presence of Symbiodiniaceae (50).

### Localisation and formation of *Endozoicomonas* aggregates in *A. loripes* tissues

In this study, we demonstrated that distinct *Endozoicomonas* phylotypes were found as aggregates within the gastrodermis of *A. loripes*. These findings align with several previous studies on other coral species (29,38,41). Most CAMAs observed in this study were located within the mesenteries of the gastrovascular cavity, consistent with the localisation of CAMAs in other acroporid corals, such as *Acropora hyacinthus* (41) and *Pocillopora acuta* (Maire et al., 2024). However, unlike these corals, CAMAs were not observed in the mesenterial filaments, specialized extensions of the mesenteries that can be extruded from the polyp’s mouth during digestion or defense (Fig. 4.). Moreover, unlike pocilloporid corals where CAMAs validated as *Endozoicomonas* are mainly detected in the epidermis of the tentacles (28,32,38), no CAMAs were detected within the tentacles of *A. loripes*. These findings suggest the existence of coral species-specific niches for CAMA formation.

Coral-associated bacteria have been reported to differ in chemotactic capabilities used to find optimal niche microhabitats within their host (51,52). Therefore, differences in CAMA location could be linked to the chemotactic capabilities of natural populations of coral-associated bacteria towards chemicals released by corals and/or their symbionts or localisation near preferred metabolites including carbohydrates, ammonium and dimethyl sulfoniopropionate (53,54). Therefore, variations in biochemical and nutritional requirements across different *Endozoicomonas* strains may impact the selection of their preferred ecological niche within the coral holobiont.

No CAMAs were found in the epidermis nor the gastrodermis of the tentacles in this study. Instead, all CAMAs co-localised with Symbiodiniaceae in gastrodermal tissues, including the actinopharynx, mesenteries, and coenenchyme. This spatial distribution suggests metabolic interactions between CAMAs and Symbiodiniaceae, where oxygenic photosynthesis generates high oxygen levels during the day. Microhabitats within the mesenteries or gastrovascular cavity may optimize microbial metabolism, with CAMAs positioned in areas with lower oxygen or temporary anoxic conditions during nighttime respiration, supported by localized high nutrient gradients such as ammonium, nitrate, nitrite, and phosphate, vitamin B_12_. In contrast, CAMAs in the epidermis, as reported in pocilloporid corals (Bayer et al., 2013; Wada et al., 2022; Maire et al., 2023), may experience lower nutrient availability in surrounding reef waters, particularly during tentacle extension for feeding (55). These differences suggest that variations in biochemical and nutritional requirements among *Endozoicomonas* strains influence their tissue localization and preferred ecological niches.

### *Endozoicomonas* composition within aggregates

CAMAs were predominantly formed by cells from the same phylogenetic clade (Clade-A or Clade-B), with a minor proportion (2%) containing the two distantly related *Endozoicomonas* phylotypes (94% average 16S rRNA sequence identity) coexisting within the same aggregate. Only one other recent study has applied species-specific probes to investigate the composition of the bacterial community residing within CAMAs of corals, in this case in *S. pistillata* (28). Their findings indicated that CAMAs were predominantly located in the tentacles and contained a single phylotype of *Endozoicomonas* or a mix of highly similar phylotypes (sequence identity ≥97.7%). Similarly, only closely related *Endozoicomonas* strains (16S rRNA V5-V6 sequence identity 96 to >99%) were excised from the same CAMA in the coral *P. acuta* (32).

Different bacterial phylotypes within aggregates can interact antagonistically through secretion systems and compete for nutrients, resulting in growth inhibition of one of the phylotypes (60,61). However, microorganisms can also take advantage of the proximity to interacting cells via bidirectional beneficial relationships (62–64). Such synergistic interactions can lead to the evolution of advantageous traits, such as metabolic complementation (65), which arises when one or multiple species utilise metabolites produced by neighbouring species. Moreover, mixed biofilms tend to exhibit enhanced resilience to host immune responses compared to their mono-species counterparts (66–68). All these interactions are often facilitated by cross-species communication via quorum sensing (69–71). Hence, the CAMA development of one *Endozoicomonas* phylotype could play an important role in governing the sequential recruitment of either conspecific strains or other *Endozoicomonas* species. Conversely, the dominance of members from Clade-B (Fig. 4), as well as the low proportion of mixed CAMAs in this study compared to *S. pistillata* (28) could suggest a degree of niche exclusion between phylogenetically distinct *Endozoicomonas* species. This is corroborated by the finding that mixed CAMAs were only obtained from one geographical region in (28).

### Aggregate morphological variation

An intriguing finding in this study was the observed differences in CAMA morphology between the two identified clades. While Clade-A exhibited CAMAs with regular and contained growth patterns, Clade-B aggregates lacked a clear boundary and exhibited an irregular shape. While the specific bacterial taxa forming CAMAs remained unknown, a previous study documented cellular-level variations among collective CAMAs within the same *A. hyacinthus* coral, including CAMAs comprised of rod-shaped, pleomorphic, filamentous-like, or spore-like structures (41). However, the morphology of bacterial cells within individual CAMAs in *A. loripes* appeared homogenous. Therefore, to our knowledge, this is the first time such systematic variations have been documented, hence this phenomenon merits further investigation. SEM confirmed aggregates in genotype Al07 were surrounded by a membrane of unknown origin (Fig. 8.). This membrane-like structure formed a clear boundary around the aggregates, potentially indicating compartmentalisation within the host tissue. While SEM and FISH could not be performed on the same CAMAs, the Al07 genotype exhibited a higher relative abundance of Clade-A *Endozoicomonas* compared to Clade-B *Endozoicomonas* (Fig. 1). This observation suggests a potential association between Clade-A *Endozoicomonas* and the membrane-like structures, possibly reflecting an intracellular localisation. However, SEM images of Clade-B CAMAs could not be obtained, leaving their ultrastructural characteristics unresolved in this study.

**Fig. 7.**
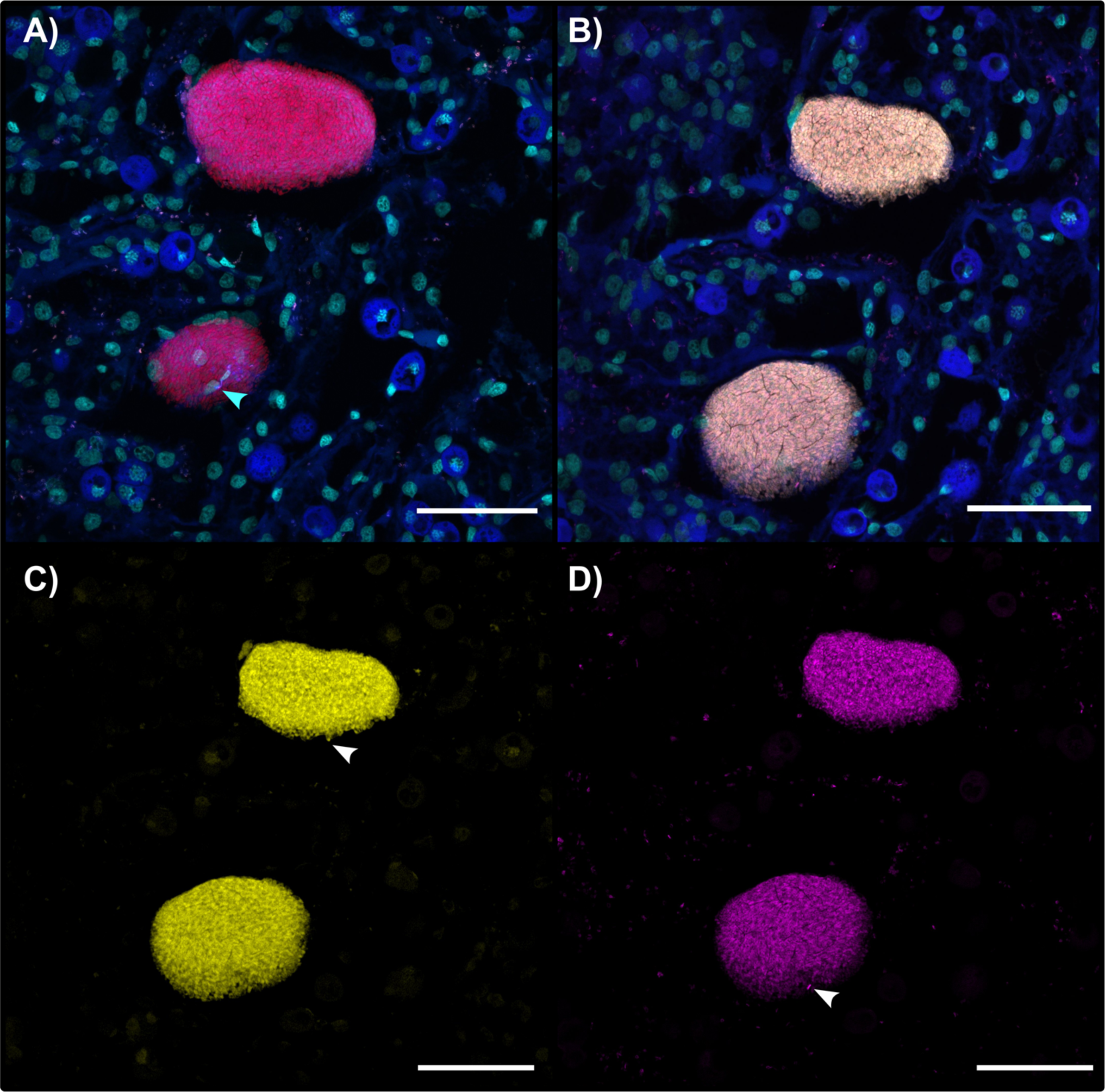
Detection and co-localisation of mixed-CAMA populations. Fluorescent signals from CAMAs binding both phylotype-specific probes. **(A)** Two adjacent CAMAs in a tissue section hybridised with the ‘All-*Endozoicomonas* mix’ probe (red) and stained with DAPI depicted in cyan. Cyan arrow indicates coral host nuclei at the periphery of a CAMA, suggesting coral cells are overlapping the aggregate. **(B)** The consecutive section to the one shown in (A) hybridised with Clade-A (magenta) and Clade-B (yellow) specific probes, showing binding of both probes within the same CAMAs. Arrows indicate individual bacterial cells hybridised by the Clade-B probe (top CAMA) and Clade-A probe (bottom CAMA) at the periphery of the CAMAs while it is dikicult to observe individual cells at the centre of the CAMA as both signals are manifested in the merged image (B-C). Coral tissue in (A) is sectioned 8 μm above (B). Blue: host autofluorescence. Cyan: nuclear structures stained by DAPI. C) and D) are split channels from the merged image shown in **B**. White arrows indicating individual cells which can be distinguished in **B**. Scale bars represent 20 μm.

**Fig. 8.**
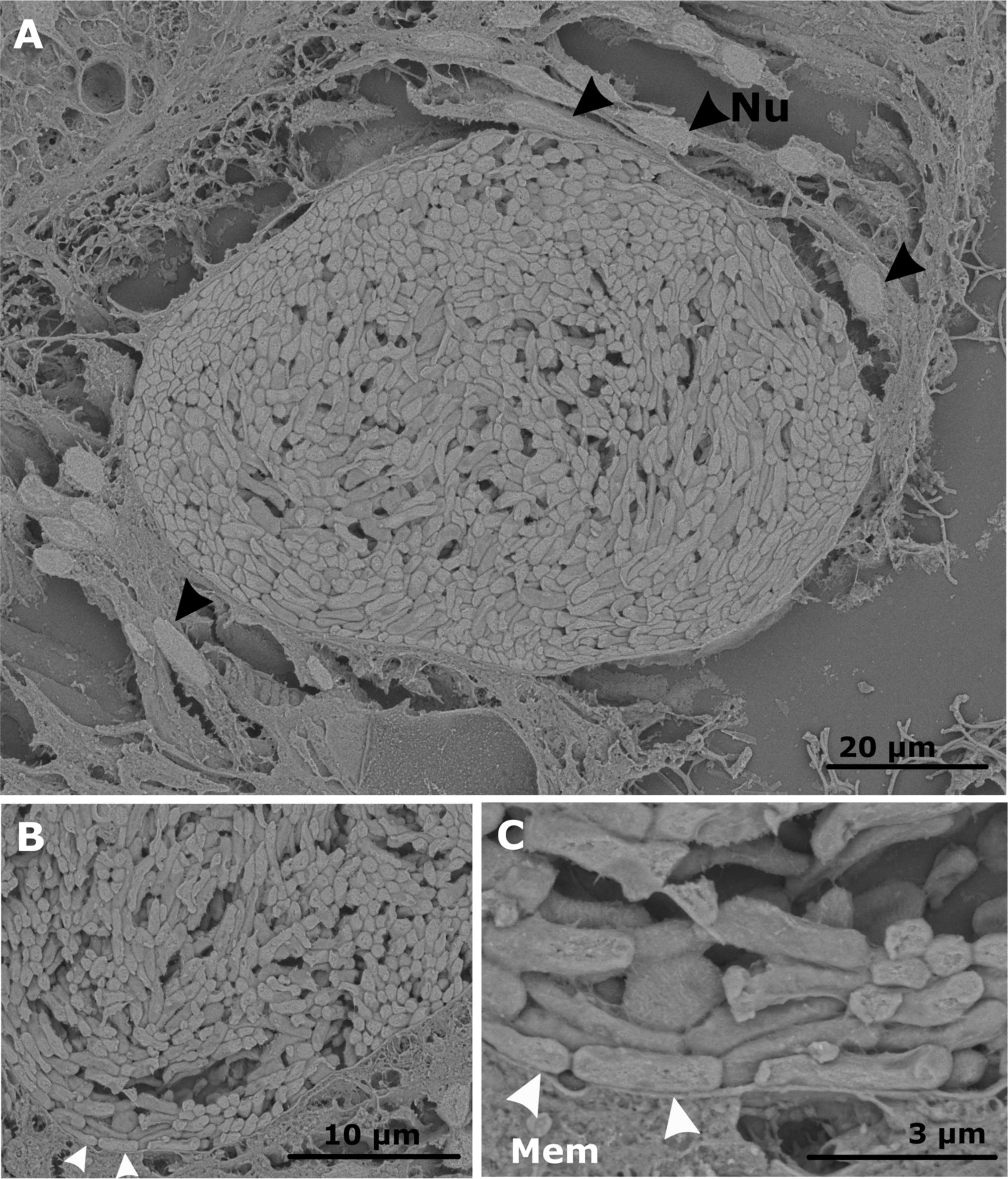
Scanning electron micrographs visualising bacterial aggregates within the coral tissues of coral genotype Al07 using back scattered electrons. **A** Overall view of a restricted bacterial aggregate within the coral tissue. Host nuclei are marked with black arrows, indicating their spatial relationship adjacent to the aggregate. **B-C** Close-up views highlighting a membrane-like structure of unknown origin surrounding the bacterial aggregate (indicated by white arrows).

The possible membranes enclosing Clade-A CAMAs imply a controlled process of aggregation, possibly involving specific adhesion molecules, such as fimbriae or targeted secretion systems (72,73). In certain cases, secretion systems of both pathogenic and symbiotic bacteria can deliver effector proteins that interact directly with host cell machinery. These effectors can facilitate adherence, promote cell invasion via host immune suppression and support the intracellular survival of bacteria within host cells (74–76). Furthermore, the regular and contained growth pattern of these aggregates within host tissues could also suggest the presence of a membrane. Notably, the presence of a membrane of unknown origin was recently observed in CAMAs within the tentacles of the coral *P. acuta* (32) suggesting an intracellular localisation. Moreover, CAMAs of unknown composition detected in the coral *Porites compressa* were found to be contained within a membrane (29). These observations also extend to the cnidarian model *Exaiptasia diaphana*, where the presence of a membrane surrounding CAMAs found within the tentacles was dependent on the size of the aggregates, with only aggregates smaller than anemone cells appearing to be intracellular (77).

Conversely, the absence of a clear margin in CAMAs formed by members of Clade-B CAMAs suggests they may utilise different adhesion molecules or extracellular matrix components, leading to a distinct pattern of aggregation within the coral tissue. Clade-B may have alternative strategies for evading host immune system cues compared with Clade-A, with consequences for subsequent internalisation or lack thereof (72,78). Moreover, the differentiating CAMA morphologies could also be reflected in metabolic differences between clades. A recent study has demonstrated that bacterial populations of marine Vibrionaceae strains secrete enzymes required for polysaccharide breakdown, which may be reflected in cell aggregation patterns (79). Specifically, strains with lower levels of lyases tend to aggregate more strongly than those that secrete high levels of enzymes. Furthermore, the study suggests that increased aggregation enhances intercellular synergy within tightly packed aggregates compared to strains that aggregate more diffusely. These findings suggest that metabolic functions can be reflected in the cell aggregation patterns of marine bacteria capable of extracellularly catabolising polysaccharides.

Lastly, the irregular and dispersed formation patterns observed for Clade-B CAMAs could also indicate a dynamic state, suggesting that these structures at any moment could be either in the process of being broken down or represent early stages of colonisation within the coral tissues. Preliminary evidence points to the fact corals can feed on their symbiotic algae by digestion of excess symbiont cells (80), raising the possibility that a similar process could be occurring in bacterial CAMAs.

## Conclusion

In summary, this study provides key insights into the role of *Endozoicomonas* in coral health and functioning. Metabarcoding and culturing revealed the dominance of *Endozoicomonas* in *A. loripes* colonies, identifying two coexisting clades. FISH experiments showed clade-specific CAMAs in the coral’s gastrodermis, with distinct morphologies suggesting differing host-microbe interactions. Notably, a membrane of unknown origin was observed enclosing CAMAs dominated by Clade-A, though further validation is required to confirm this structure. Moreover, the genomic potential of these clades and mechanisms of CAMA formation will be explored in future studies.

## Acknowledgements

We acknowledge the Wulgurukaba and Bindal peoples as the first scientists and traditional custodians of the lands and sea country where this research was conducted. We extend our sincere gratitude to the SeaSim team for their collaboration and support in coral collection and husbandry. We are also grateful to Chien Lit Cheah for assistance with histology and staining. Additionally, appreciate the Biological Optical Microscopy Platform (The University of Melbourne) for confocal microscopy training and support with image analysis, the BioSciences Microscopy Unit and the Ian Holmes Imaging Centre for optical and electron microscopy.

## Data Availability Statement

The datasets generated during the presented study are available under NCBI BioProject ID PRJNA1198616.

## Authors’ contributions

Conceptualization: C.R.G., L.H., L.L.B., and M.J.H.v.O. Formal analysis: C.R.G., Funding acquisition: M.J.H.v.O and L.L.B. Investigation: C.R.G., A.D., A.v.d.M., K.D. and G.K.P. Methodology: C.R.G., A.D., L.H., L.L.B., and M.J.H.v.O. Supervision: L.L.B., L.H. and M.J.H.v.O. Visualization: C.R.G., and J.M., Writing—original draft preparation: C.R.G., L.H., L.L.B., and M.J.H.v.O.

