## Supplementary materials for "Differential aggregation patterns of *Endozoicomonas* within tissues of the coral *Acropora loripes*"

### Supplementary methods

#### Coral collection and maintenance

Visually healthy *A. loripes* colonies (i.e., showing no signs of tissue necrosis or bleaching) with a diameter averaging ~25 cm were collected at 3-6 m depth from two sites on the central Great Barrier Reef. At Davies Reef (18°51”S; 147°63”E), five colonies were collected in June 2020 and four were collected in February 2021. At Back Numbers Reef (18°31'23.4”S; 147°08'39.9”E), five colonies were collected in December 2020 (all at 3-6 m depth). Colonies were transported to the Australian Institute of Marine Science (AIMS, Townsville) in aerated 80-L plastic bins receiving a steady flow of seawater. Once at AIMS, the colonies were transferred into 1200-L tanks in the outdoor experiment area of the National Sea Simulator (SeaSim), where they were supplied with 0.4 µm filter-sterilised seawater (FSW) (top-up rate 4 L $min^{-1}$, 4.8 turnovers per day) and fed daily with *Artemia* at 0.5 nauplii $mL^{-1}$. Corals were kept under ambient lighting and temperature conditions following the Davies Reef profile.

#### 16S rRNA gene and ITS2 metabarcoding of coral samples

To characterise the bacterial composition of *A. loripes* collected from the field, coral fragments were sampled from four different locations within each coral colony to capture as much microbial diversity as possible. This approach was chosen due to the significant variation in microbial communities both within and among colonies of *A. loripes*, as previously observed by (19). These 4 replicate samples were taken across four within-coral locations ranging from shaded at the bottom of colonies to sun exposed areas at the apical tip of coral branches. To minimise cross-contamination, nitrile gloves were discarded between each sampling location, and all collection equipment (bone cutters and forceps) was sequentially sterilised in 10% sodium hypochlorite, RO water, 80% ethanol and lastly a wash in 0.22 µm FSW. Genomic DNA extractions were carried out on 3-5 mm replicate coral fragments, followed by targeted amplification of the variable regions V5-V6 of the bacterial 16S rRNA gene and the ITS2 region as described previously (19). Sequencing of the obtained amplicons was carried out at the Walter and Eliza Hall Institute (Melbourne, Australia). Subsequent processing of 16S rRNA gene amplicon sequences was carried out in QIIME2, with statistical analyses performed in R in accordance with (20). Raw ITS2 sequences were submitted to SymPortal for ITS2 profiling (81) and subsequent SymPortal analysis were carried out as detailed previously (82). For a complete list of PCR primers used in this study refer to Table S1.

#### Bacterial culturing

For bacterial isolation, 3 cm sized coral fragments were sampled using sterile bone cutters and forceps and rinsed in 0.22 µm FSW (Whatman) to remove loosely associated microorganisms. Fragments were then placed in separate sterile zip-lock bags containing 10 ml autoclaved filtered, 0.22 µm FSW. Using an air gun with a sterilised nozzle, tissue was sprayed off the skeleton and the slurry was transferred into 50-mL polypropylene conical tubes and homogenised for 30 s at 3000 rpm using a tissue homogeniser (HoverLabs, India) as described for the targeted isolation of tissue-associated *Endozoicomonas* (83). The resulting tissue homogenate was then plated in triplicate onto Marine Agar 2216 (MA; BD Difco) in the following dilutions (1:10, 1:100, 1:1000, 1:10,000). Bacterial growth started to appear after four days of incubation at 23°C in the dark and bacterial colonies were purified from a single bacterial colony after a minimum of three clean passages onto fresh MA plates. For the establishment of bacterial stock cultures, pure bacterial colonies were transferred into individual tubes containing 5 mL Marine Broth 2216 (MB; BD Difco) and incubated at 170 rpm at 28°C for 48 h in a temperature-controlled incubated shaker (New Brunswick Innova 40, Eppendorf). Cells were harvested from the mid-exponential phase and aliquots of the bacterial culture were prepared as 30% glycerol stocks, snap-frozen and subsequently cryopreserved at -80 °C.

Individual bacterial colonies were screened by colony PCR amplification per (84) with minor modifications. Briefly, bacterial colonies were stabbed with a sterile pipette tip, suspended in 20 µL of lysis solution (0.05 M NaOH, 0.25% SDS), and heated for 15 min at 95 °C, followed by centrifugation at 4000 × *g* for 5 min to obtain genomic DNA in the supernatant. A volume of 1 µL supernatant was dispensed into a PCR tube containing 20 µl of PCR master mix (1x AmpliTaq Gold PCR Master Mix, 1 µM of the *Endozoicomonas*-specific primer En771R (85), and 1 µM of the bacterial universal primer 27F (86)). Amplification was carried out under the following conditions: 94°C for 3 min, followed by 35 cycles of each: 30 s at 94°C, 30 s at 55°C, 45 s at 72°C followed by a final extension step set at 72°C for 10 min. Bacterial colonies that amplified with the *Endozoicomonas*-specific primer were then subjected to new PCR reactions using the universal bacterial primer pair 27F/1492R (86). Post-PCR clean-up and Sanger sequencing using the 1492R primer were carried out at Macrogen Inc (Seoul, South Korea). Reverse complement reads from each isolate were subjected to quality control by trimming low-quality bases and aligned using the global alignment tool in Geneious Prime 2021.2.2 and poor-quality base pairs were trimmed using the Trim Ends tool. The 16S rRNA gene sequences from each isolate were also used for homology search against the National Centre of Biotechnology Information (NCBI) non-redundant database using Basic Local Alignment Search Tool (BLAST) (GenBank)(87).

#### Phylogenetic relationships within the genus *Endozoicomonas*

To assess how well the cultured strains represented the *Endozoicomonas* communities associated with the coral fragments, a custom BLAST database encompassing all ASVs from the metabarcoding data was created using Geneious Prime 2021.2.2. Subsequently, sanger sequences from each cultured isolate were queried against this database and a relative abundance plot of 100% sequence identity matches was generated. To determine the phylogenetic placement of the *Endozoicomonas* strains isolated from *A. loripes*, the 16S rRNA sequences obtained from Sanger sequencing were used to construct a phylogenetic tree with near full-length Endozoicomonadaceae 16S rRNA sequences retrieved from the SILVA database (88). The sequences were aligned with MAFFT for SSU rRNA alignment (89), with a total alignment length of 2087bp. A maximum likelihood (ML) phylogenetic tree was inferred using IQ-TREE v1.6.11 (90). ModelFinder was utilised within IQ-TREE to automatically select the most suitable model (91). The TIM3e+R6 model was chosen based on Bayesian Information Criterion values with 1000 bootstraps to examine the statistical robustness of the tree topology. Finally, the interactive Tree of Life iTOL was used to draw the final consensus tree (92).

#### Design of *Endozoicomonas* phylotype-specific probes

Sequences of the 16S rRNA gene for each isolate were aligned and a consensus sequence for both Clade-A and Clade-B was constructed using the global alignment tool in Geneious Prime 2019.1.2 (https://www.geneious.com). The consensus sequence length for each clade ranged from averaged around 1200 nucleotides. These consensus sequences were selected as target sequences for designing FISH probes.

Two phylotype-specific oligonucleotide probes unique to Clade-A and Clade-B were created using the ARB probe design tool (93) with a custom reference sequence database. This database comprised a total of 362,515 prokaryotic 16S rRNA gene sequences of cultured bacteria from the ARB/SILVA SSU Ref dataset release v. 138.1 (88). Additionally, it incorporated >1500 16S rRNA gene sequences of *Endozoicomonas* spp. sourced from publicly available databases. To identify conserved and variable regions suitable for probe design, a multiple-sequence alignment of all rRNA gene sequences was performed using the global SINA Aligner (v1.2.12).

The parameters GC content, melting temperature ($T_m$), and secondary structure formation were considered to optimise probe performance. Additionally, BLAST searches were performed against public databases to ensure probe specificity, and probes showing potential cross-reactivity with non-target sequences were further refined. Lastly, the required stringency of the hybridisation conditions and formamide concentrations were evaluated *in silico* using mathFISH against increasing formamide gradients (94).

#### Testing probes in ARB against *A. loripes* metabarcoding sequences

To ensure the robustness of FISH experiments, probes were designed to minimise the number of matches between each clade and with other non-target sequences. While the probe targeting Clade-B had 4 mismatches with Clade-A, the probe targeting Clade-A had 2 mismatches with members of Clade-B (Fig. 1). However, designing clade-specific probes with enough mismatches between them to ensure specificity, as well as having zero non-target hits was unattainable. As both metabarcoding and FISH samples were collected simultaneously, sequences identified as potential non-target hits in ARB were extracted and compared against a custom database in Geneious, which comprised all amplicon sequence variants obtained from 16S metabarcoding. This allowed us to estimate which non-target organisms were present within our samples (Fig. S2). The relative abundance of non-target hits in the amplicon data was plotted in RStudio using ggplot2 (95).

From this analysis, we selected a Clade-A-specific probe with the fewest potential mismatches, despite sharing 100% sequence similarity with three non-target ASVs in the metabarcoding data. Similarly, for the Clade-B-specific probe, only one non-target hit was identified, sharing 100% similarity with an ASV in the metabarcoding data. Since all FISH experiments utilized both clade-specific FISH probes and an 'All-*Endozoicomonas* mix' probe, competitor probes were exclusively designed against non-target groups within the *Endozoicomonas* genus (Table S2-2). This approach ensured that non-specific binding with any non-*Endozoicomonas* cells could be verified by the absence of signal from the 'All-*Endozoicomonas* mix' probe. The specificity of the probes was confirmed by BLASTn searches against the NCBI database NCBI). The finalised oligonucleotide probe sequences for Clade-A and Clade-B were modified at the 5' end with either Atto647N or Atto550 and synthesised by Biomers (Ulm, Germany).

#### Fluorescence *in situ* hybridization of bacterial cell suspensions

To evaluate the stringency of each probe (i.e., Endo-Clade-A and Endo-Clade-B), FISH was carried out in solution on cultured pure isolates using a series of formamide concentrations for each probe (Fig.3). Each isolate was grown in Marine Broth (MB) medium (DIFCO 2216) (28°C, 180 rpm), with cells harvested at late logarithmic phase (~48 h after inoculation) and fixed in 4% (v/v) paraformaldehyde for 4 h at 4℃. Prior to in-solution FISH, cells were concentrated by centrifugation and washed twice in PBS. Aggregated clumps of *Endozoicomonas* cells were separated using a sonicator for 2 × 15 s cycles on high (Power Sonic 505, Thermoline Scientific)

Cells pelleted by centrifugation (5000 rpm, 5 min) were resuspended in 100 µl FISH hybridisation buffer (0.9 M NaCl, 20 mM Tris-HCl pH 7.4, 1% SDS) with increasing formamide concentrations (15 - 30% formamide with 5% increments) and incubated with 5 ng $\mu l^{-1}$ final concentration of each respective probe (Biomers, Germany) at 46°C for 2 h. Following hybridisation, pelleted cells were washed twice in 100 µl of pre-warmed (48°C) wash buffer (20 mM Tris-HCl, 5 mM EDTA, 0.01% SDS, 0.080 M NaCl) followed by incubation at 48 °C for 20 min. Finally, cells were centrifuged twice in 100 µl of ice-cold 1x PBS and resuspended in ultra-pure water. Optical microscope filters were selected according to the fluorochromes used (max. excitation/emission in nm: Atto550 485/498; Atto647N 490/525). The bacterial cells were imaged at 40X magnification using a confocal laser scanning microscope LSM890 (Zeiss) With Zen2.3 software (Black) from Zeiss. The *in vitro* probe evaluation yielded two Clade-specific probes, which only hybridised with their intended target, i.e., Probe-A hybridised only with isolates from Clade-A, and Probe-B exclusively hybridised with Clade-B isolates (Fig. S3). Furthermore, both the Endo-Clade-A (Fig. S4) and Endo-Clade-B (Fig. S5) probes demonstrated high stringency *in situ*. This was evidenced by minimal background staining with only non-specific binding directed at nematocytes and mucocytes, which is a well-known artefact of *in situ* detection of bacterial assemblages within coral tissues (Fig. S4)(96).

#### Sample fixation and histological processing

During sample collection for metabarcoding, A. *loripes* fragments from the same coral colonies were fixed and preserved for histology and FISH. Five replicate 1-2 cm-sized fragments were cut from five different locations of each coral colony as described above. Each fragment was immediately fixed for 24 h in 4% paraformaldehyde (ProScitech, AUS) at 4°C, rinsed in phosphate-buffered saline (PBS) and preserved in 50% ethanol (in PBS) at -20°C until further processing. Fixed coral fragments were rinsed twice in PBS (20 min each) and subsequently decalcified in 10% ethylenediaminetetraacetic acid (EDTA-$Na_2 \cdot 2H_2O$, Sigma-Aldrich, USA; adjusted to pH 8) in a rotary tube mixer at 4°C for approximately 3 weeks. The EDTA solution was changed every 2 days until no skeleton was left and the tissue appeared completely clear (14–60 days depending on the fragment size of each sample). Decalcified tissue samples were rinsed in PBS and dehydrated sequentially in 70%, 80%, 90% and 100% ethanol (40 min each), followed by 3 washes in 100% ethanol (60 min each) and 3 immersions in xylene (60 min each), before embedding in paraffin wax (Paraplast Plus, Fisher Scientific, USA), with all steps performed using an automated vacuum infiltration processor (Tissue-Tek® VIP, Sakura, Japan). Five replicate sections from all coral colonies were cut longitudinally (in the median plane of the polyp mouth) and horizontally (in the transverse plane of the polyp mouth), sectioned at 4 μm and mounted on either glass slides for H&E staining or Superfrost® slides for FISH experiments.

#### Hematoxylin and Eosin (H&E) staining

Before staining, each serial tissue section was dewaxed in xylene (3 x 30 s), and rehydrated by passing tissue sections through decreasing concentrations of ethanol 100% (3 x 30 s), 70% (30 s) and lastly phosphate-buffered saline (PBS). Hydrated sections were stained in Mayer's hematoxylin (3 min), washed in water, then dipped in Scott's tap water until sections became blue, followed by tap water wash and lastly counterstained with Eosin Y and further rinsed in tap water. Following staining, sections were dehydrated in reverse order of ethanol 70% (30 s), 100% (90 s), cleared in xylene (90 s) and lastly mounted in ProLong Antifade mountant (ThermoFisher). H&E slides were examined with a Zeiss AxioImager M2 microscope using Zen Blue (Zeiss, Germany). The "area" tool from the software ImageJ (97) was used to measure CAMA surface areas on H&E slides in Fiji (98).

#### FISH on coral tissue

Whole mount FISH with *Endozoicomonas*-specific probe was carried out as described previously (32). FISH on tissue sections were conducted according to established protocols with minor adjustments (99). Briefly, each serial tissue section was dewaxed in xylene (2x 10 min), after which semi dried slides were passed through increasing concentrations of methanol to quench host tissue autofluorescence (50% for 5 min, 75% for 5 min, 90% for 5 min, 100% methanol for 5 min), immediately followed by three washes in 100% ethanol (3x 5 min). Then tissue sections were permeabilised with hydrochloric acid for 12 min, and lastly rinsed in 20 mM Tris-HCl solution (pH 8.0) for 10 min and air-dried. Probe hybridisation was carried out with an adjusted probe concentration of 5 ng $\mu l^{-1}$. Following hybridisation, slides were counterstained with the nuclear stain DAPI (final concentration 5 µg $\mu l^{-1}$; Merck, Germany) and mounted with cover glass using Citiflour antifadent mountant (proSciTech, Australia). For each coral genotype, five replicate sections from five branches were observed.

#### FISH Microscopic Analysis

The 'All-*Endozoicomonas* mix' probe and Clade-A probe were labelled with Atto647N, while the non-sense probe and Endo-Group-B probe were labelled with Atto550. Each experiment involved probing two consecutive sections: one with the 'All-*Endozoicomonas* mix' and non-sense probe and one with Clade-A and Clade-B specific probes. Probes were detected with the following laser settings: 405 nm (0.4%), 488 nm (0.2%), 561 nm (0.06%), and 633 nm (0.07%). Due to highly autofluorescent host tissue, the entire emission spectra of both host pigments (tissue section without probe) and FISH-stained bacterial isolates were recorded using spectral scanning on a confocal laser scanning microscope LSM890 (Zeiss) With Zen2.3 software (Black) from Zeiss. Linear unmixing of the obtained emission spectra was conducted to separate host tissue from the signal emitted by the FISH probes. For a complete list of FISH probes used in this study refer to Table S2.

#### Scanning electron microscopy

Histology sections of the ALOR7 sample were dewaxed and stained with haematoxylin and eosin stain followed by imaging using a Leica DM6000 widefield optical microscope (Leica Microsystems). After detection of CAMAs, the same sections were then heavy metal stained for scanning electron microscopy (SEM) using 1% uranyl acetate for 10 min followed by lead citrate for a further 2 min. After washing, sections were dried and carbon coated before imaging on a

Hitachi SU7000 FE SEM (Hitachi) at 3 kV. Images were acquired simultaneously using the upper detector for high resolution secondary electrons, the mid detector for in-lens back scatter electrons or the lower detector for secondary electrons. The CAMA were located via SEM via determination of the optical microscope map and final images were processed using FIJI and Inkscape.

#### Supplementary Figures

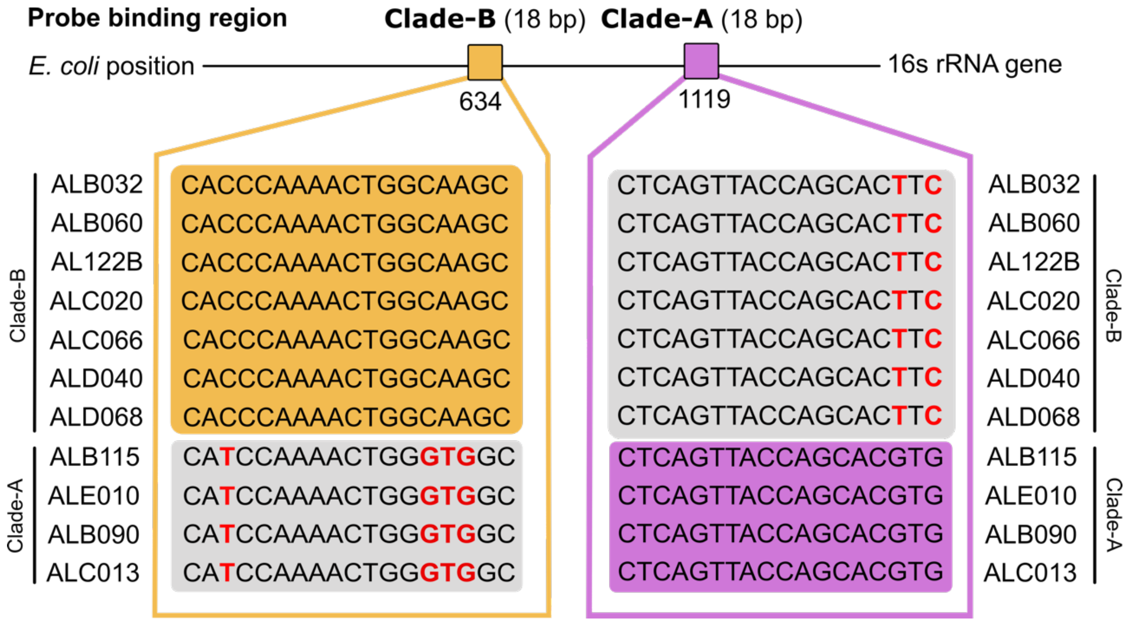


**Fig. S1** Alignment of 16S rRNA Sanger sequences used for FISH probe design. The Clade-A and Clade-B probes are designed to specifically target their respective clades, with the Endo-Clade A probe showing four mismatches with Clade-B members and the Endo-Clade B probe showing two mismatches with Clade-A members at the binding sites. Mismatches are highlighted in red.

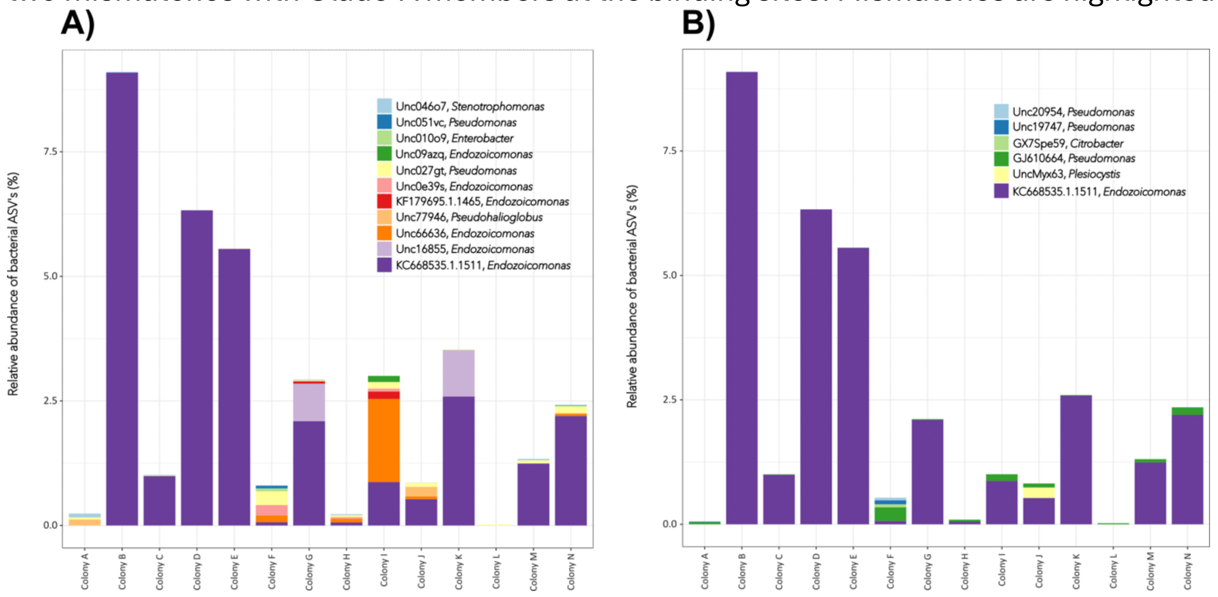


**Fig. S2.** Relative abundance of non-target probe hits present in the metabarcoding data. **A** Displays all potential non-target hits of Endo-Clade-A against sequences found within the tissues of *A. loripes* **B** Displays all potential non-target hits of Endo-Clade-B against sequences found within the tissues of *A. loripes*. Competitor probes targeted Unc0e39s, KF179695.1.1465, Unc66636, Unc16855, KC668535.1.1511 blocking the potential non-target probe binding site was created for Endo-Clade A. Likewise, a competitor probe targeted KC668535.1.1511 blocking the potential non-target probe binding site was created for Endo-Clade

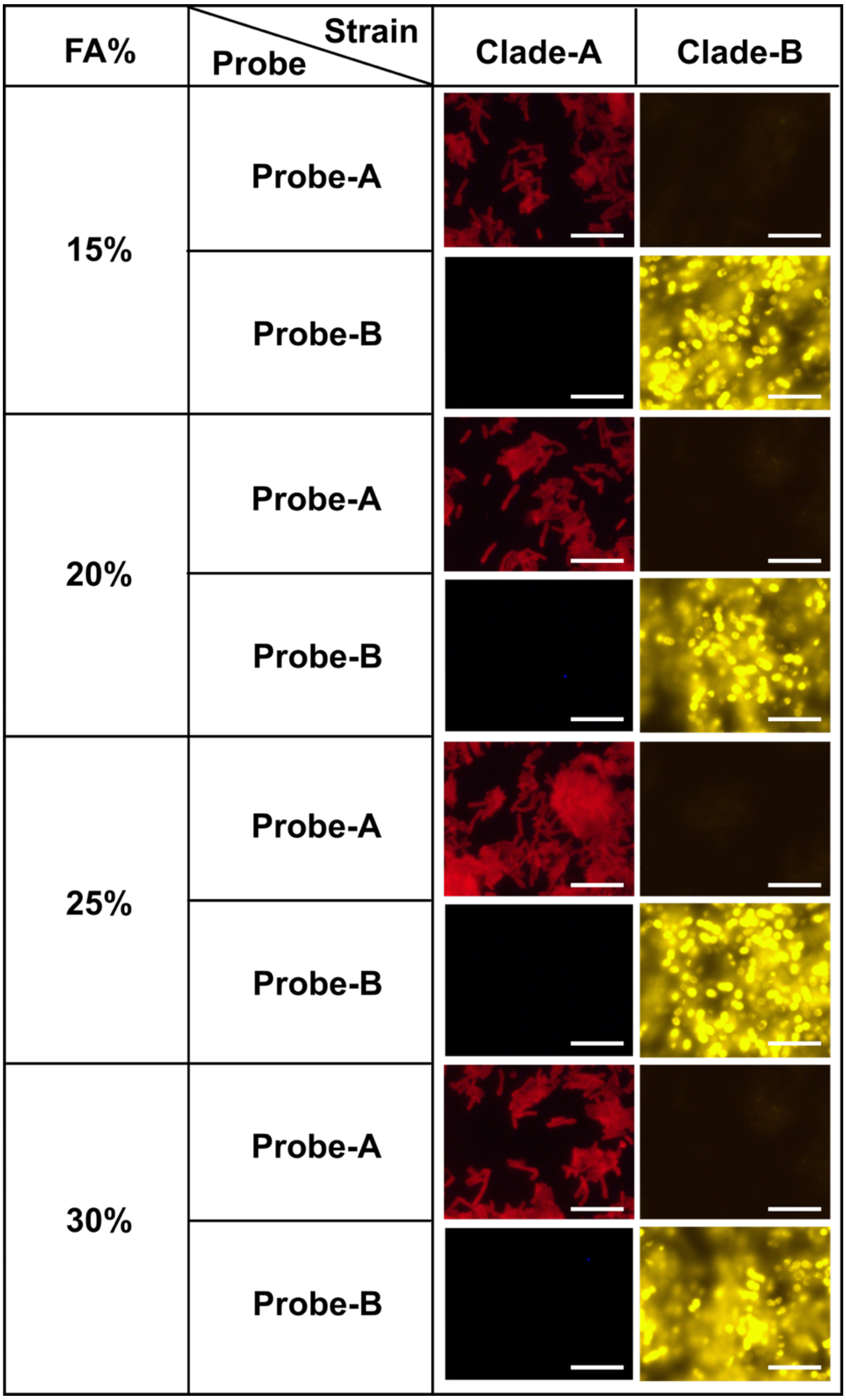


**Fig. S3.** *In vitro* probe evaluation. Microphotographs of bacterial isolates from Clade-A and Clade-B hybridised with each respective probe. *In vitro* testing of the Endo-Clade A probe and Clade-B probe on bacterial isolates cultured from Clade-A and Clade-B respectively revealed highly specific binding patterns. No evidence of probe hybridization was observed in non-target groups, indicating precise targeting and strong discriminatory capability against members of Clade-A and Clade-B. These results also affirmed stringency across all formamide concentrations (FA%). Scale bars represent 5 μm.

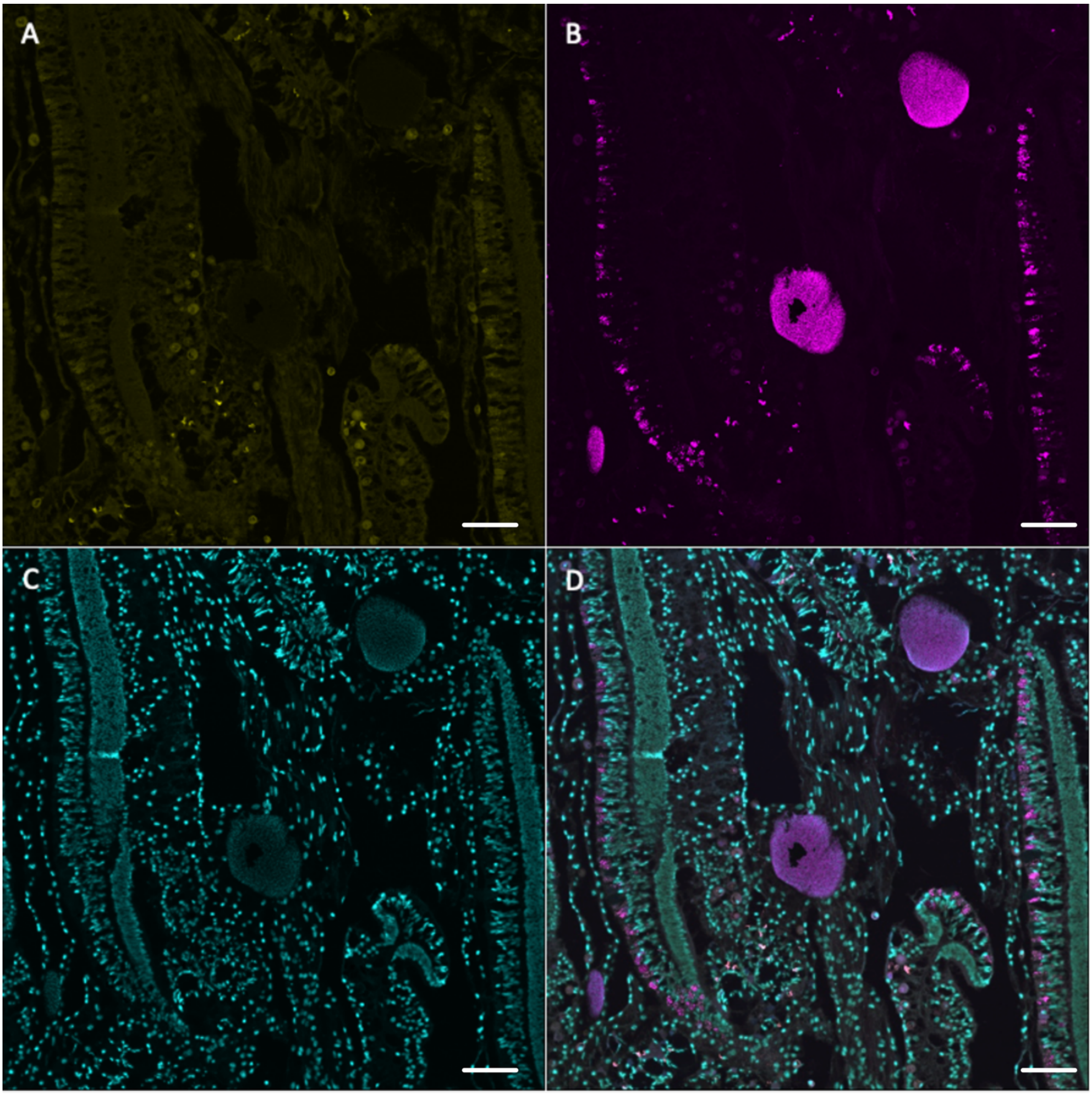


**Fig. S4.** Fluorescence *in situ* hybridization images showing specific probe binding within a CAMA from *A. loripes* from coral genotype Al08. **A** Displays no binding of Endo-Clade B probe **B** shows binding by Endo-Clade A probe. **C** Nuclear structures stained by DAPI **D** Merged image of all signals. Scale bars represent 100 µm.

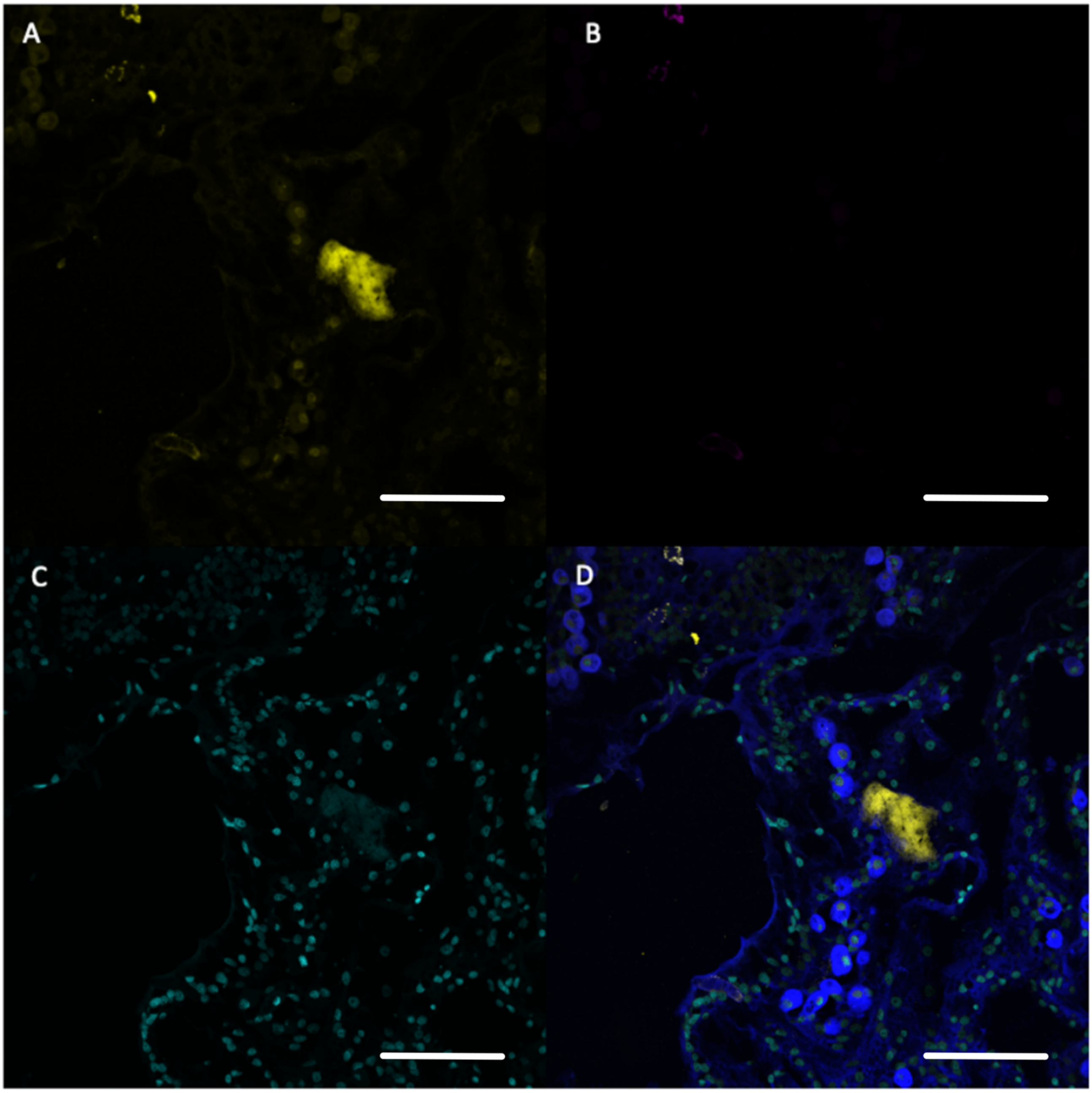


**Fig. S5** Fluorescence *in situ* hybridization images showing specific probe binding within a CAMA from *A. loripes* genotype Al08**. A** Shows binding by Endo-Clade B probe. **B** Displays no binding of Endo-Clade A probe **C** Cellular structures stained by DAPI **D** Merged image of all signals. Scalebars represent 20 µm.

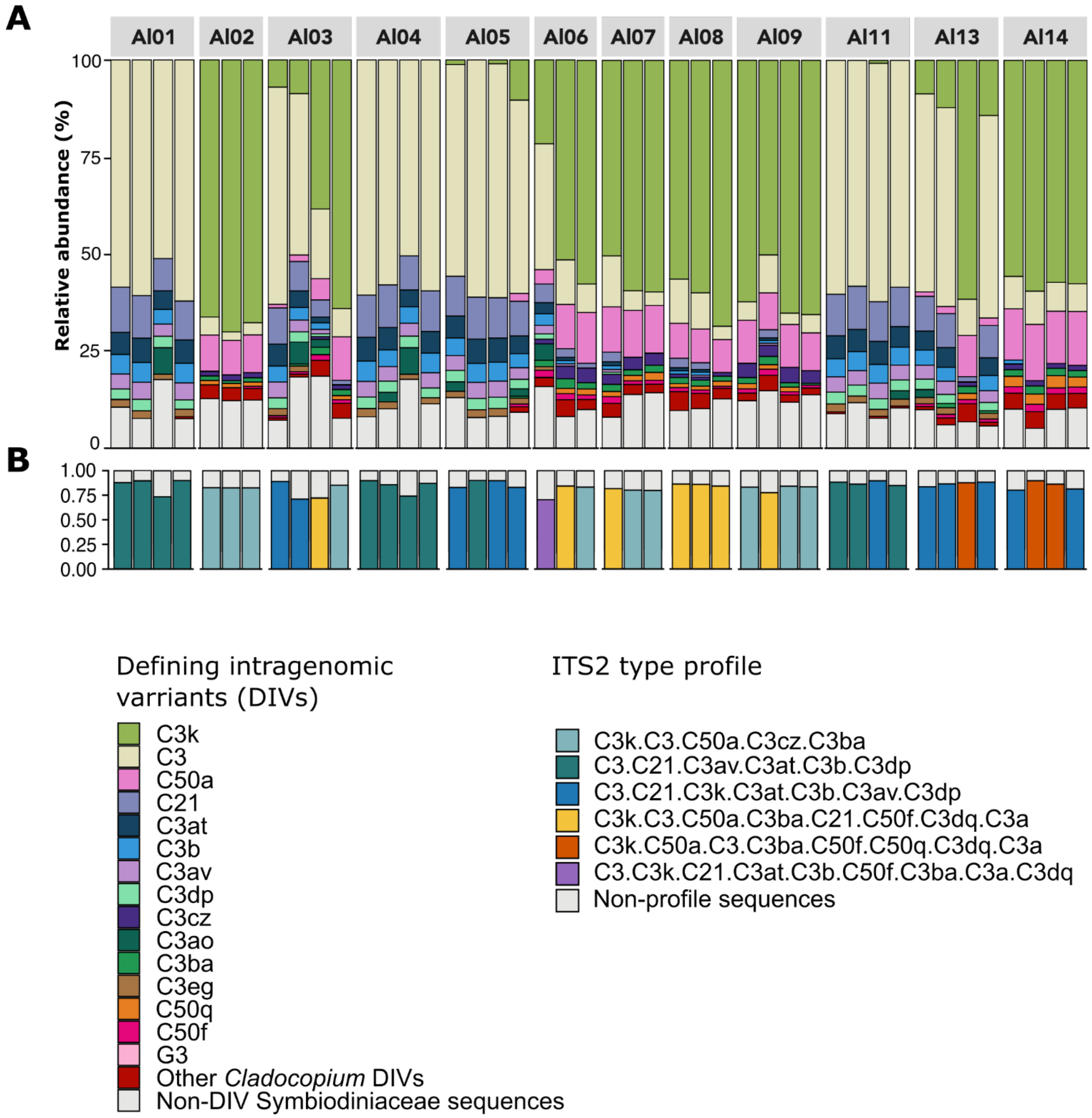


**Fig. S6** Relative abundances of ITS2 sequences and predicted ITS2 type profiles. **A** The stacked bar charts display the relative abundances of defining intragenomic variants (DIVs), while **B** shows the corresponding ITS2 type profiles. Each bar represents a replicate sample, with four replicates sequenced per coral colony, as indicated on the X-axis. Colours denote the relative abundance of DIVs **A** or ITS2 profiles **B**. Only the top 15 major ITS2 sequences are shown, while the remaining sequences are grouped under "other *Cladocopium* DIVs" and "non-DIV symbiodiniaceae sequences". Sequences were processed using the SymPortal analytical framework to classify DIVs and generate ITS2 type profiles.

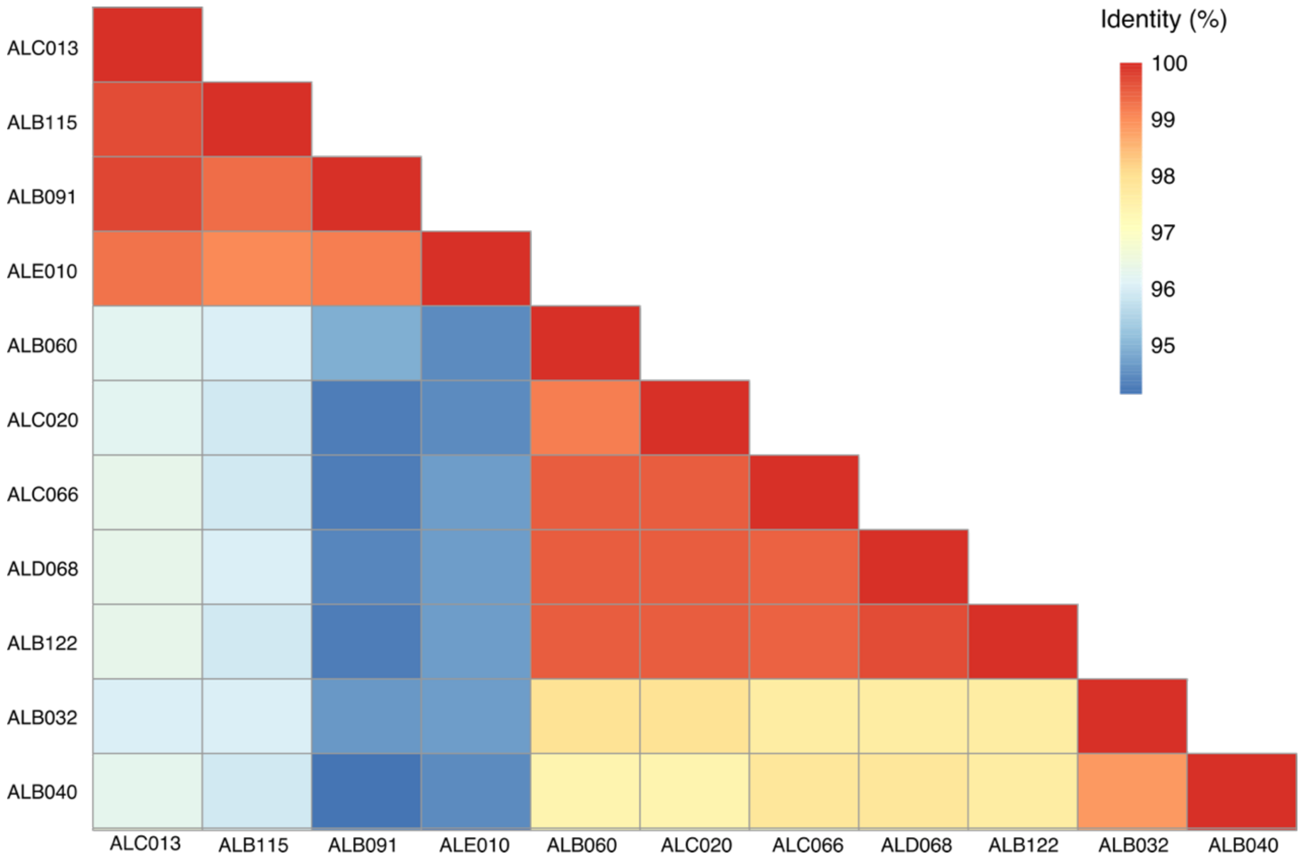


**Fig. S7** Strain similarities based on rRNA gene sequence similarity matrix calculated from pair-wise comparisons. Six strains showed high sequence identity (ALB060, ALC020, ALC066, ALD068, ALB122, ALB032, ALB040) among them, while they have a lower sequence identity of approximately 94% with the remaining 4 strains (ALC013, ALB091, ALB115, ALC013) revealing clear species boundaries of < 95% ANI.

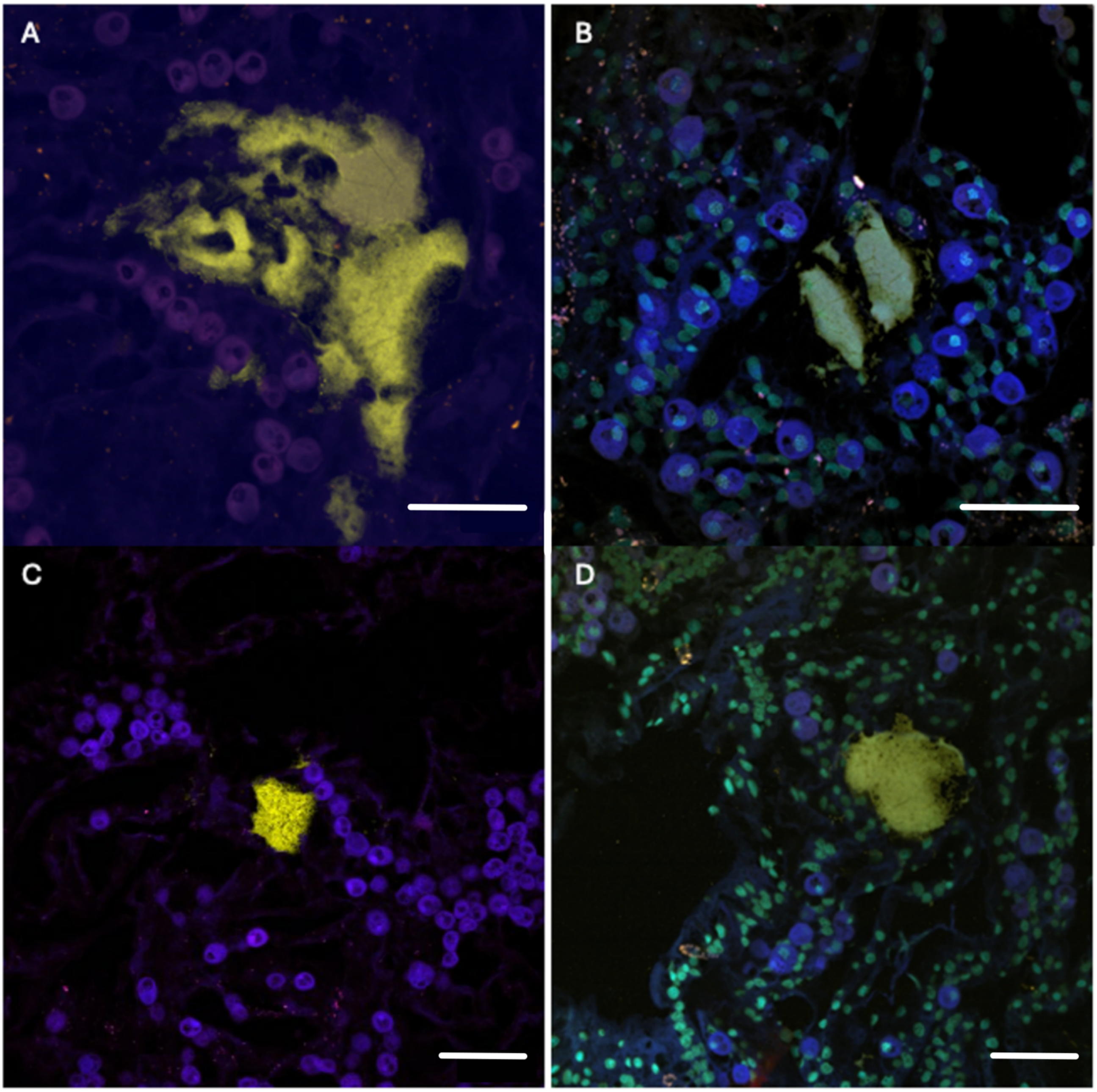


**Fig. S8** FISH images showing Clade-B CAMA morphotype spectrum**.** Clade-B CAMA morphotypes range from highly dispersed aggregation patterns **A-B** to somewhat more circular shapes **C-D**, however, no clear boundary appears in the outer periphery. **A-D** Yellow: Clade-B specific-probe. Red: Non-EUB probe (negative control). Blue: Host autofluorescence. Orange: overlap between target probe and negative control probe indicating non-specific binding with nematocytes and mucocytes. **B** & **D** Cyan: Host nuclei stained by DAPI. Scalebars represent 20 µm.

#### Supplementary tables

**Table S1** List of primers used in this study

| DNA region | Primer name | Forward/ reverse | Primer sequence 5' - 3' | Reference |
|---|---|---|---|---|
| 16S rRNA gene | 784F | Forward | TCGTCGGCAGCGTCAGATGTGTATAAGAGACAG<br>AGGATTAGATACCCTGGTA | (100) |
| 16S rRNA gene | 1061R | Reverse | GTCTCGTGGGCTCGGAGATGTGTATAAGAGACAG<br>CRRCACGAGCTGACGAC | (100) |
| ITS2 | SYM_VAR_5.8S2 | Forward | GTGACCTATGAACTCAGGAGTC<br>GAATTGCAGAACTCCGTGAACC | (101) |
| ITS2 | SYM_VAR_REV | Reverse | CTGAGACTTGCACATCGCAGC<br>CGGGTTCWCTTGTYTGACTTCATGC | (101) |
| 16S rRNA gene | 1492R | Forward | AGAGTTTGATCMTGGCTCAG | (86) |
| 16S rRNA gene | 27F | Reverse | TACGGYTACCTTGTTACGACTT | (86) |
| 16S rRNA gene | En771R | Reverse | TCAGTGTCARRCCTGAGTGT | (85) |

**Table S2** List of all oligonucleotide probes used for Fluorescence *in situ* Hybridization.

| Target group | Probe | Sequence (5'-3') | Formamide % | Reference |
|---|---|---|---|---|
| All bacteria | EUB338-mix | GCWGCCWCCCGTAGGWGT | 35 | (40) |
| Negative control | nonEUB | ACATCCTACGGGAGG | 35 | (102) |
| Endozoicomondaceae | Endozoi663 | AGGAGUGUGGAAUUUCC | 35 | (38) |
| Endozoicomondaceae | Endozoi736 | CUCUGGUCUGACACUGAC | 35 | 38) |
| Clade A | Endo-Clade-A | CACGTGCTGGTAACTGAG | 35 | This study |
| Clade B | Endo-Clade-B | CACCCAAAACTGGCAAGC | 35 | This study |
| Clade A competitor | Com-Endo-CladeA1 | CACGTGCTGGTAACTAAG | 35 | This study |
| Clade A competitor | Com-Endo-CladeA2 | TACGTGCTGGTAACTGAG | 35 | This study |
| Clade A competitor | Com-Endo-CladeA3 | GAAGTGCTGGTAACTGAG | 35 | This study |
| Clade B competitor | Com-Endo-CladeB | GCTTGCCAGTTTTGGATG | 35 | This study |
